# Elucidation of tumor-stromal heterogeneity and the ligand-receptor interactome by single cell transcriptomics in real-world pancreatic cancer biopsies

**DOI:** 10.1101/2020.07.28.225813

**Authors:** Jaewon J. Lee, Vincent Bernard, Alexander Semaan, Maria E. Monberg, Jonathan Huang, Bret M. Stephens, Daniel Lin, Brian R. Weston, Manoop S. Bhutani, Cara L. Haymaker, Chantale Bernatchez, Cullen M. Taniguchi, Anirban Maitra, Paola A. Guerrero

**Affiliations:** Sheikh Ahmed Center for Pancreatic Cancer Research, The University of Texas MD Anderson Cancer Center, Houston, TX, USA; Department of Translational Molecular Pathology, The University of Texas MD Anderson Cancer Center, Houston, TX, USA; Department of Surgical Oncology, The University of Texas MD Anderson Cancer Center, Houston, TX, USA; Department of Radiation Oncology, The University of Texas MD Anderson Cancer Center, Houston, TX, USA; Department of Experimental Radiation Oncology, The University of Texas MD Anderson Cancer Center, Houston, TX, USA; Department of Gastroenterology, Hepatology, and Nutrition, The University of Texas MD Anderson Cancer Center, Houston, TX, USA; Department of Melanoma Medical Oncology, The University of Texas MD Anderson Cancer Center, Houston, TX, USA; Department of Pathology, The University of Texas MD Anderson Cancer Center, Houston, TX, USA

**Keywords:** pancreatic ductal adenocarcinoma, single-cell RNA sequencing, cancer biopsy, molecular subtypes, tumor microenvironment

## Abstract

Precision medicine approaches in pancreatic ductal adenocarcinoma (PDAC) are imperative for improving disease outcomes. However, the long-term fidelity of recently deployed *ex vivo* preclinical platforms, such as patient-derived organoids (PDOs) remains unknown. Through single-cell RNA sequencing (scRNA-seq), we identify substantial transcriptomic evolution of PDOs propagated from the parental tumor, which may alter predicted drug sensitivity. In contrast, scRNA-seq is readily applicable to limited biopsies from human primary and metastatic PDAC and identifies most cancers as being an admixture of previously described epithelial transcriptomic subtypes. Integrative analyses of our data provide an in-depth characterization of the heterogeneity within the tumor microenvironment, including cancer-associated fibroblast (CAF) subclasses, and predicts for a multitude of ligand-receptor interactions, revealing potential targets for immunotherapy approaches. While PDOs continue to enable prospective therapeutic prediction, our analysis also demonstrates the complementarity of using orthogonal *de novo* biopsies from PDAC patients paired with scRNA-seq to inform clinical decision-making.

**Statement of Significance:** The application of single-cell RNA sequencing to diagnostic pancreatic cancer biopsies provides in-depth transcriptomic characterization of the tumor epithelium and microenvironment, while minimizing potential artifacts introduced by an intervening *ex vivo* passaging step. Thus, this approach can complement the use of patient-derived organoids in implementing precision oncology.

## Introduction

The majority of patients with pancreatic ductal adenocarcinoma (PDAC) present with locally advanced or metastatic disease, which precludes surgical resection of their cancer (1). The gold standard for diagnosis is a tissue biopsy prior to initiation of systemic therapy, which is obtained by either endoscopic ultrasound-guided fine needle aspiration (EUS-FNA) of the primary lesion or through interventional radiology (IR)-guided percutaneous biopsy of a metastatic site. While typically adequate for histopathological assessment, PDAC biopsies, especially those obtained via EUS-FNA without rapid onsite cytopathology assessment, can be limited by low neoplastic cellularity (2,3). Nonetheless, such biopsies may be the only source of tissue from patients with locally advanced or metastatic PDAC who are likely to derive the most benefit from tailored approaches.

Multiple recent studies have demonstrated the feasibility of using next generation sequencing (NGS) platforms for mutational analysis as applied to these limited “real world” biospecimens obtained from PDAC patients (4,5). Notably, although targeted NGS panels can identify somatic or germline DNA mutations in a clinically meaningful timeframe, no more than a quarter of PDAC patients, at best, harbor such actionable mutations (6). Therefore, efforts at expanding precision medicine approaches in PDAC beyond DNA alterations have led to patient-derived organoids (PDOs) being used for prospective therapeutic prediction. While initial reports primarily utilized surgical resection samples for establishing PDOs, more recently, the feasibility of using limited biopsy material has also been confirmed, garnering the possibility of incorporating PDOs into precision medicine trials in patients with advanced PDAC (7,8). Nonetheless, although PDOs are a facile preclinical platform for therapeutic prediction, how their molecular landscape might evolve during passaging, compared to the parental tumor, is less well established. This is undoubtedly pertinent, since transcriptomic shifts occurring *ex vivo* may impact sensitivity or resistance to predicted agents, with implications for how this information may be used for clinical decision-making.

In this study, we characterize the transcriptomic landscape of PDOs in the course of *ex vivo* passaging, and the potential impact on therapeutic prediction. Using single-cell RNA sequencing (scRNA-seq), we find that *ex vivo* culture of PDOs leads to transcriptomic shifts that are distinct from the parental tumor, as well as from earlier passages of a given PDO. We then explore the feasibility of using scRNA-seq on *de novo* limited biopsies obtained from patients with primary and metastatic PDAC. As recently described in studies using surgically resected PDAC samples, scRNA-seq provides an unprecedented level of insight into the architecture of the neoplastic cells and their tumor microenvironment (TME), including compartment specific heterogeneity (9–11). We demonstrate that scRNA-seq on limited endoscopic and core biopsies captured nearly all of the previously reported repertoire of cell types in surgical resections, including the tumor-stromal heterogeneity inherent to this disease, and revealed putative mechanisms for immune evasion within the TME. We believe that precision medicine approaches to a near universally lethal disease like PDAC should deploy multiple orthogonal approaches, including NGS for actionable DNA alterations, PDOs for phenotypic therapeutic prediction, and scRNA-seq on contemporaneous *de novo* biopsies for assessment of the tumor-stromal architecture. In particular, scRNA-seq on *de novo* biopsies could potentially minimize the confounding effects of transcriptomic shifts in *ex vivo* passaged PDOs on therapeutic predictions.

## Results

### *Ex vivo* Propagation of Patient-Derived Organoids Leads to Transcriptomic Shifts

PDOs have emerged as an important tool for prospective therapeutic prediction across multiple cancer types (7,8,12–15), but whether they retain the molecular architecture of the parental tumor following *ex vivo* propagation is unknown. We first performed scRNA-seq on a PDAC PDO (PDO-1) after 5 and 15 *ex vivo* passages to examine whether the transcriptomic machinery evolves in culture over time. We found that in later passage organoids, there were two transcriptomically distinct clusters (**Supplementary Figure S1A**), which formed two separate cell fates on trajectory inference (**Supplementary Figure S1B**). Transcripts upregulated towards cell fate 1 showed enrichment of epithelial-mesenchymal transition (EMT), reactive oxygen species (ROS) biosynthesis, response to growth factor, and axon development (**Supplementary Figure S1C**). In comparison, transcripts upregulated towards cell fate 2 were enriched for telomere maintenance, DNA replication, cell cycle, and hypoxia pathways (**Supplementary Figure S1C**). Copy number inference from scRNA-seq revealed 16p gain in late-passage organoids and 19q loss specifically in cell fate 2 (**Supplementary Figure S1D**). Our results suggest that *in vitro* culture conditions shift the transcriptomic landscape towards deregulated proliferative pathways and those indicative of aggressive phenotypes, such as EMT and hypoxia.

To identify transcriptomic changes from *in vivo* parental tumor to *ex vivo* culture, we obtained a different PDO sample (PDO-2) from a PDAC patient with vaginal apex metastasis, a biopsy from which was used for *de novo* scRNA-seq and PDO generation. After 14 passages, at which time there was an adequate number of cells to perform additional studies, we performed scRNA-seq to compare to the parental tumor sample. Epithelial cells from the tumor and organoids formed distinct clusters (**Figure 1A**), and gene set enrichment analysis (GSEA) revealed the enrichment of cell cycle-related pathways such as E2F targets, G2M checkpoint, and mitotic spindle in PDO-2. In contrast, the parental tumor was enriched for immune-related pathways such as interferon (IFN) α and γ signaling, tumor necrosis factor (TNF) α signaling, as well as hypoxia pathway (**Figure 1B**). Trajectory inference analysis showed an unbranched progression from tumor to PDO-2, and dynamic gene expression changes along the trajectory once again demonstrated downregulation of IFN-related pathways and upregulation of cell cycle pathways with pseudotime (**Figures 1C and 1D**). A prior scRNA-seq analysis of non-passaged surgically resected samples also found the upregulation of inflammation-related pathways in PDAC epithelial cells (10), suggesting that these signals likely arise as a result of cues from the TME *in vivo* and are lost during organoid culture.

**Figure 1.**
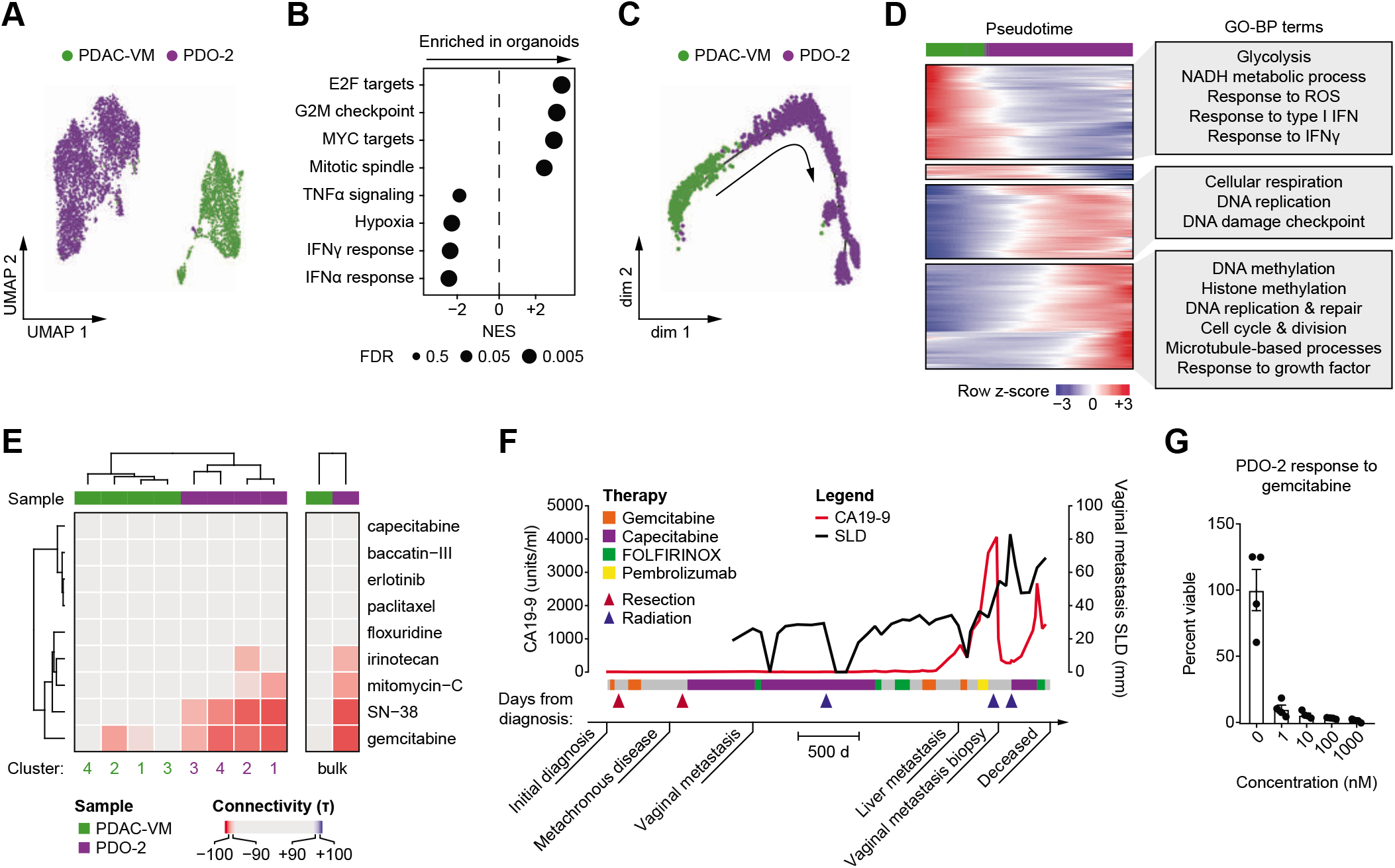
Patient-derived organoids evolve with time. **A.** Uniform manifold approximation and projection (UMAP) plot of single cells from epithelial compartment of PDAC vaginal metastasis tissue (PDAC-VM, green) and organoids after 14 passages (PDO-2, purple). **B.** Bubble plot showing enrichment of pathways in organoids compared to tissue with normalized enrichment score (NES) on the x-axis. Size of the bubble represents false discovery rate (FDR). **C.** Pseudotime trajectory of single cells from PDAC-VM (green) and PDO-2 (purple). **D.** Heatmap showing dynamic gene expression changes through pseudotime (left) and enrichment of Gene Ontology biological processes (GO-BP) terms for each gene cluster (right). **E.** Heatmap showing connectivity score for transcriptomically distinct clusters from PDAC-VM (green) and PDO-2 (purple) to commonly used compounds in PDAC (left) and the same for pseudo-bulk samples (right). Negative connectivity score predicts sensitivity. **F.** Clinical course of vaginal metastasis patient. CA19-9 is plotted with red line and sum of longest diameter (SLD) of the target lesion is plotted with black line. Significant events and treatment courses are displayed on the x-axis. **G.** Bar graph showing cell survival of PDO-2 after treatment with gemcitabine at different concentrations. Error bars represent standard error of mean (S.E.M.).

We also tested whether there were copy number alterations (CNAs) in addition to transcriptomic changes in the transition from tumor tissue to PDOs. We performed both single-cell copy number sequencing (**Supplementary Figure S1E**) and CNA inference from scRNA-seq profiles (**Supplementary Figure S1F**). In the parental neoplastic cells, both methods showed large copy number events across multiple chromosomes. In PDOs, there were additional CNAs and more heterogeneity in copy number profiles. These results once again demonstrate that long-term maintenance of PDOs may lead to both transcriptomic and genomic alterations.

To assess whether these transcriptomic changes led to an altered response to therapy, we performed drug sensitivity prediction using the next generation connectivity map (16). When interrogating transcriptomically distinct sub-clusters (**Supplementary Figures S1G and S1H**) and pseudo-bulk expression profiles, the cultured PDO cells were predicted to have greater sensitivity to commonly used chemotherapeutic agents in PDAC, including gemcitabine, compared to parental neoplastic cells (**Figure 1E**). Consistent with the parental neoplastic cells not demonstrating a predicted sensitivity profile to gemcitabine, the patient was clinically deemed resistant to gemcitabine prior to biopsy (**Figure 1F**); in contrast, PDO-2 was exquisitely sensitive (**Figure 1G**). Overall, based on the connectivity analysis, neoplastic cells from the parental tumor were predicted to be sensitive to 65 compounds, compared to 131 compounds in PDO-2, with a set of 50 compounds that were overlapping (“invariant”) between the parental tumor and the PDO (**Supplementary Figure S1I**).

Taken together, these data suggest that, in the course of *ex vivo* propagation, PDOs undergo transcriptomic shifts and CNAs, relative to the parental tumor as well as relative to earlier passages, with possible confounding of therapeutic prediction data. Despite the divergence in transcriptomic profiles between the PDO and parental tumor, connectivity analysis on matched scRNA-seq data revealed an “invariant” subset of predicted agents that were overlapping between the parental tumor and resulting PDO (even at passage 14). This “invariant” subset of agents in common between the parental lesion and the PDO likely reflects critical and persistent vulnerabilities in the transcriptomic circuitry, and could be prioritized for preclinical testing in the resulting PDO. However, this mandates that we can conduct high-quality scRNA-seq profiling on the *de novo* non-propagated biopsies obtained contemporaneously to the samples from which we establish PDOs.

### scRNA-seq of Limited *De Novo* Biopsies Recapitulates the Tumor-Stromal Heterogeneity of PDAC Observed in Surgical Resections

In light of our aforementioned findings in *ex vivo* propagated PDOs, and the likely need for orthogonal profiling on *de novo* samples, we explored the feasibility of conducting scRNA-seq in a panel of limited “real world” biopsies obtained from primary and metastatic PDAC patients. Five primary samples (P1-P5) were obtained via EUS-FNAs of the pancreas, while four metastatic samples were obtained as core needle biopsies (liver, lung) or segmental biopsies (vaginal apex, peritoneal). Pooled analysis of all samples yielded a total of 31,720 cells organized into 8 clusters (**Figure 2A**), and highly expressed genes in each cluster were used to identify cell types (**Figure 2B**). Epithelial cells were the most commonly represented cell type (47%), followed by T cells (28%) and myeloid cells (12%; **Figure 2A**). Across primary and metastatic lesions, all cell types were represented and the average cell composition was similar (**Supplementary Figure S2A**, **Supplementary Table S1**).

**Figure 2.**
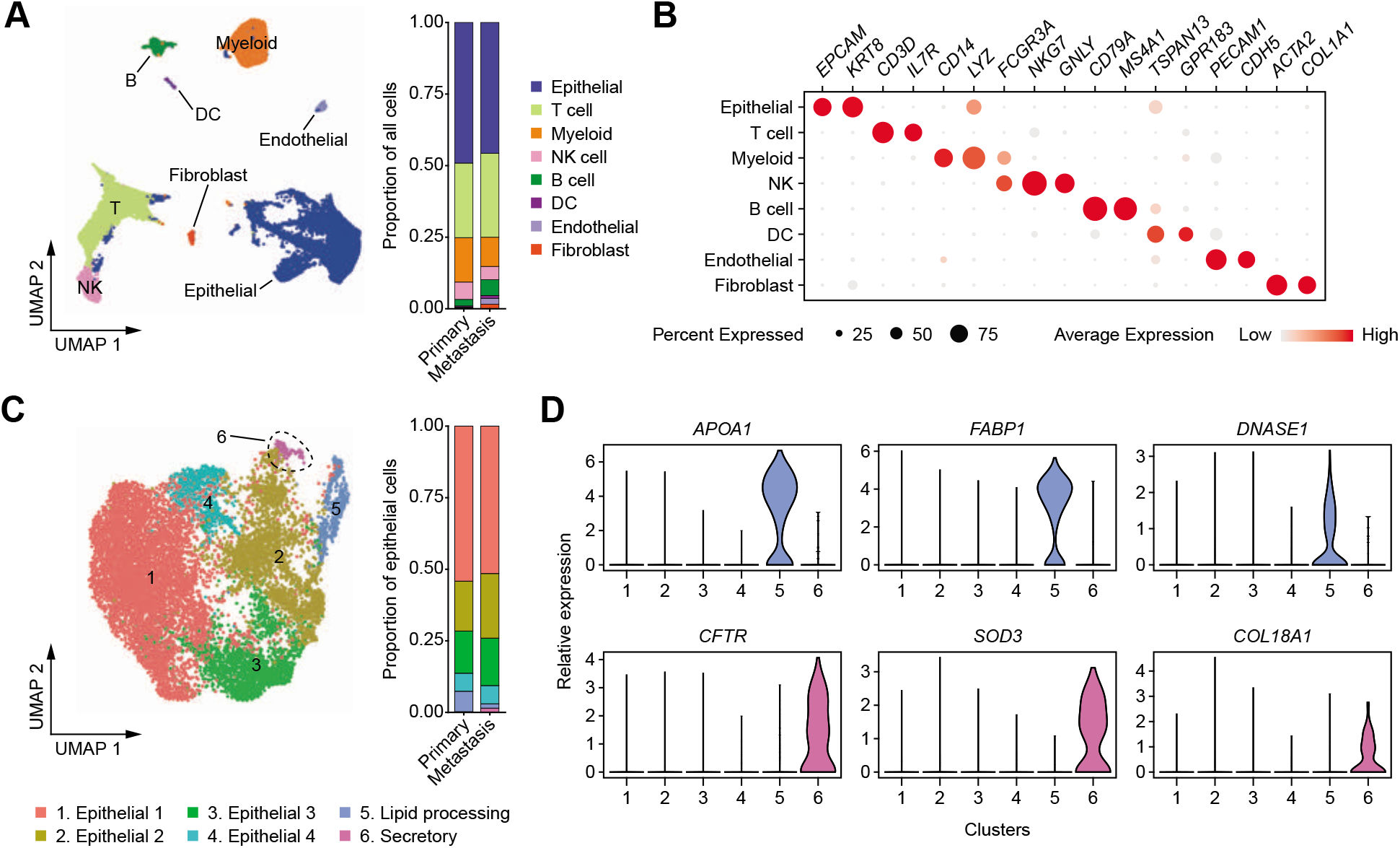
PDAC biopsies contain diverse cell types. **A.** UMAP plot of single cells from 9 biopsy samples (left) and bar plot showing proportions of cell types in pooled primary and metastatic samples (right). **B.** Bubble plot showing highly expressed marker genes in each cell type, with cell types in rows and genes in columns. Size of each bubble represents percent of cells expressing marker and color represents the level of expression. **C.** UMAP plot of epithelial cells re-clustered from A (left) and bar plot showing proportions of epithelial cell sub-clusters in pooled primary and metastatic samples (right). **D.** Violin plots showing relative expression levels of selected marker genes in all epithelial sub-clusters.

To assess the heterogeneity specific to cancer cells, we extracted the epithelial cluster for further analysis, which revealed six distinct transcriptomic sub-clusters (**Figure 2C; Supplementary Table S2**). Epithelial 1 sub-cluster was characterized by enrichment of antigen presentation, type I IFN response, and response to IFNγ pathways (**Supplementary Figure S2B**). Along the same line, epithelial 2 sub-cluster was enriched for response to ROS and intrinsic apoptosis, while epithelial sub-clusters 3 and 4 were enriched for epigenetic process and DNA replication/cell cycle, respectively, altogether demonstrating diverse transcriptomic programs present in PDAC tumor cells. A previous scRNA-seq analysis of PDAC demonstrated lipid processing and secretory cells within ductal epithelial cells (10), which were also present in our samples. Specifically, sub-cluster 5 expressed lipid processing cell markers *APOA1, FABP1*, and *DNASE1*, while sub-cluster 6 expressed secretory cell markers *CFTR, SOD3*, and *COL18A1* (**Figure 2D**). All epithelial sub-clusters were present in both primary and metastatic tissues, but lipid processing and secretory sub-clusters were proportionately less common in metastases (**Supplementary Figure S2C; Supplementary Table S2**). Copy number inference from scRNA-seq showed numerous alterations in sub-clusters 1-4, whereas lipid processing and secretory sub-clusters did not contain matching alterations (**Supplementary Figure S2D**), suggesting that the latter were derived from normal epithelium, as previously postulated (10).

In the subsequent Results, we discuss in further detail elements of tumor and stromal (including immune) diversity captured by scRNA-seq on the biopsy specimens.

#### Molecular Subtyping of Single Cells Reveals Intratumoral Subtype Heterogeneity

Previous studies have classified PDAC into molecular subtypes based on their bulk transcriptome. These classifications include Bailey subtypes (pancreatic progenitor, squamous, aberrantly differentiated endocrine exocrine or ADEX, and immunogenic), Collisson subtypes (classical, quasimesenchymal or QM, and exocrine-like), and Moffitt subtypes (classical and basal-like) (17–19). A subsequent study demonstrated two common epithelial subtypes, namely classical (or pancreatic progenitor) and basal-like (or squamous/QM), whereas immunogenic and ADEX or exocrine-like subtypes were associated with low tumor purity (20). To precisely delineate the cellular contributions to bulk molecular subtype, we used nearest template prediction (21) to classify all single cells into a subtype based on the three classifiers (**Supplementary Figures S3A-S3D; Supplementary Table S3**). Aligning the cell type to molecular subtypes confirmed that immunogenic, ADEX, and exocrine-like subtypes were mostly composed of non-epithelial cells (**Supplementary Figures S3E**), consistent with the previous finding of low tumor purity in these subtypes (20).

To understand the molecular subtypes in the context of epithelial cells, we reapplied the classifiers on the epithelial subset (**Supplementary Figures S3A-S3D and S3F; Supplementary Table S4**). Of note, while subtype prediction from pseudo-bulk projection yielded one molecular subtype for each biopsy sample (**Supplementary Figure S3G**), all tumor samples were composed of more than one subtype at single-cell level regardless of the classifier (**Figures 3A-C; Supplementary Figures S3H-S3J**), consistent with recent reports that Moffitt classical and basal-like subtypes coexist in a single tumor (22,23).

**Figure 3.**
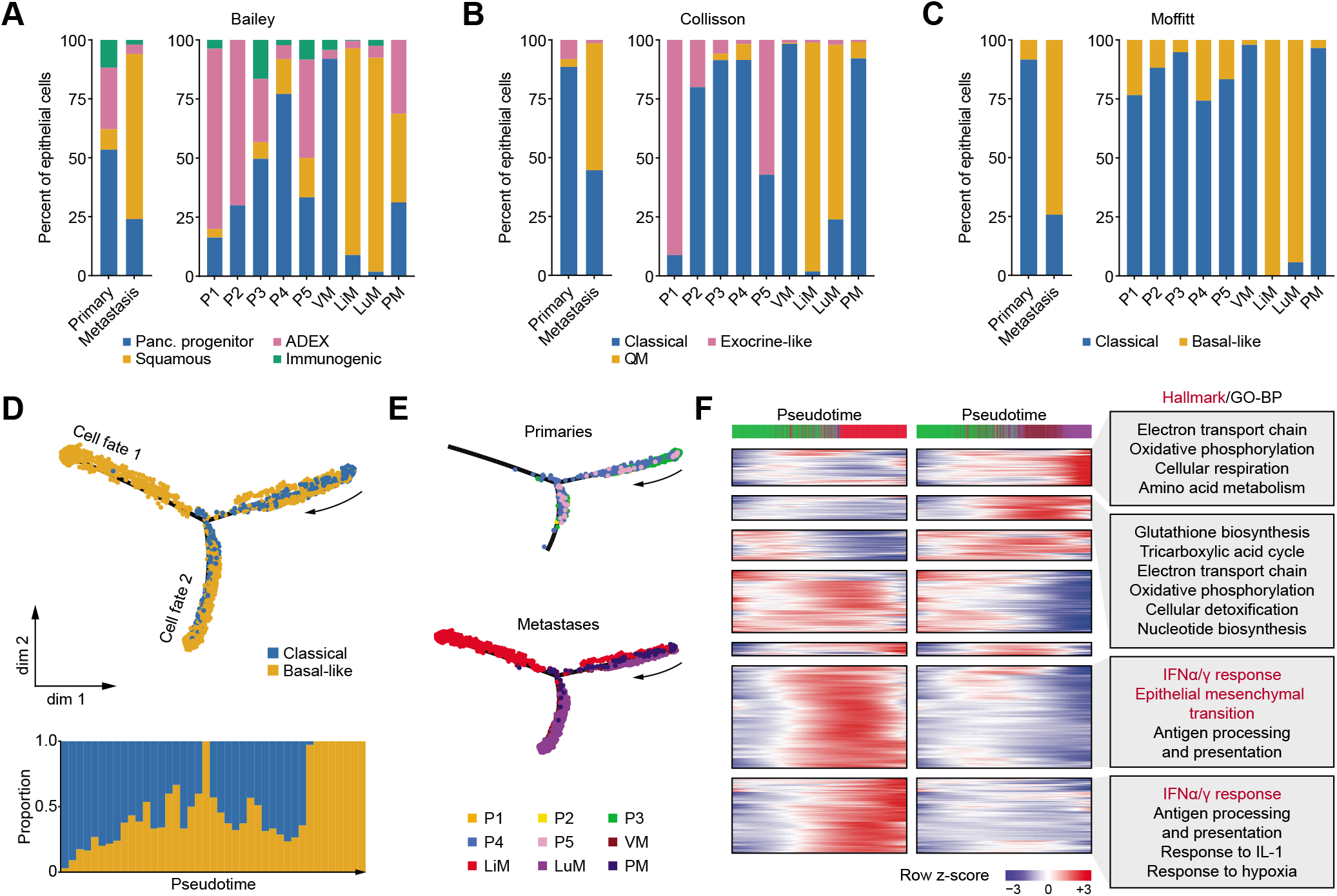
PDAC molecular subtypes at single-cell resolution. **A-C.** Bar plots showing proportions of epithelial cells classifying into Bailey (**A**), Collisson (**B**), and Moffitt (**C**) subtypes with FDR < 0.2 in pooled primary and metastatic samples (left) and individual samples (right). **D.** Pseudotime trajectory of epithelial cells from all biopsy samples labeled with Moffitt molecular subtypes (top) and bar plot representing the proportions of cells in each pseudotime bin (50 bins total) classifying into either subtype (bottom). **E.** Pseudotime trajectory plot from D labeled with sample identification separated into primary lesions (top) and metastatic lesions (bottom). **F.** Branched heatmap showing dynamic gene expression changes in epithelial cells along the pseudotime trajectory, with pseudotime progressing from left to right. Enriched Hallmark or Gene Ontology biological process (GO-BP) terms for each gene cluster are listed on the right. P1-5, primary 1-5; VM, vaginal apex metastasis; LiM, liver metastasis; LuM, lung metastasis; PM, peritoneal metastasis.

We performed pseudotime analysis on the epithelial cells to elucidate potential evolutionary trajectories of PDAC tumor cells, which revealed two major branches (**Figure 3D**). Interestingly, cells from the liver metastasis formed their own branch in the trajectory (**Figure 3E**), which may be consistent with the observation that PDAC patients with liver metastases often have poorer outcomes compared to patients with metastases in other organs including lung (24,25). Specifically, cells in the liver metastasis trajectory showed upregulation of EMT and hypoxia pathways (**Figure 3F**), which may correlate to their more aggressive phenotype. Of note, given that the liver- and lung-metastatic cells form diverging trajectories with enrichment of distinct gene sets even though cells in both sites mostly classified as basal-like (**Figures 3C and 3D**), our results suggest that there may be heterogeneity even within a single subtype that may be responsive to the metastatic site.

#### scRNA-seq on PDAC Biopsies Capture Cancer-Associated Fibroblast Diversity

In the PDAC TME, functionally and transcriptomically distinct subtypes of cancer-associated fibroblasts (CAFs) are present. Inflammatory CAFs (iCAFs) secrete cytokines and are pro-tumorigenic, whereas myofibroblastic CAFs (myCAFs) are characterized by the expression of extracellular matrix (ECM) components and exert anti-tumorigenic effects via unknown mechanisms (26). A third subtype of CAFs, named antigen-presenting CAFs or apCAFs, express class II major histocompatibility complex (MHC) genes (10). Initial analysis of the CAF population from our scRNA-seq revealed two transcriptomically distinct sub-clusters, which were designated as iCAFs and myCAFs based on the expression of *PDGFRA, CXCL12, ACTA2*, and *TAGLN* as previously described (**Supplementary Figures S4A and S4B**). To assess the presence of apCAFs, we looked for co-expression of the invariant chain of MHC II molecule *CD74* with *HLA-DPA1, HLA-DRA*, and *HLA-DRB1*, which was present in a subset of CAFs (**Figure 4A**). Utilizing nearest template prediction (21), we categorized each fibroblast into one of the three subtypes (**Figures 4B and 4C**) and found no significant difference in the proportions of the CAF subtypes in primary and metastatic tumors (**Supplementary Figure S4C; Supplementary Table S5**). In our samples, apCAFs did not co-express *PTPRC* or *MSLN* (**Supplementary Figure S3D**), confirming that these cells are not immune or mesothelial cells, as has been suggested in a recent study using murine PDAC samples (27). While apCAFs express MHC II molecules and can present antigens to CD4 T cells, they lack the co-stimulatory molecules required to activate an immune response. Therefore, it was hypothesized that apCAFs wound contribute to the immunosuppressive environment in PDAC by leading to anergy or differentiation of CD4 T cells into regulatory T (Treg) cells (10). Indeed, the proportion of apCAFs in biopsy samples was negatively correlated with the ratio of memory and effector CD8 T cells to Tregs (**Figure 4D**). Our results recapitulate a potential mechanism by which stromal cells within the PDAC microenvironment may lead to immune suppression.

**Figure 4.**
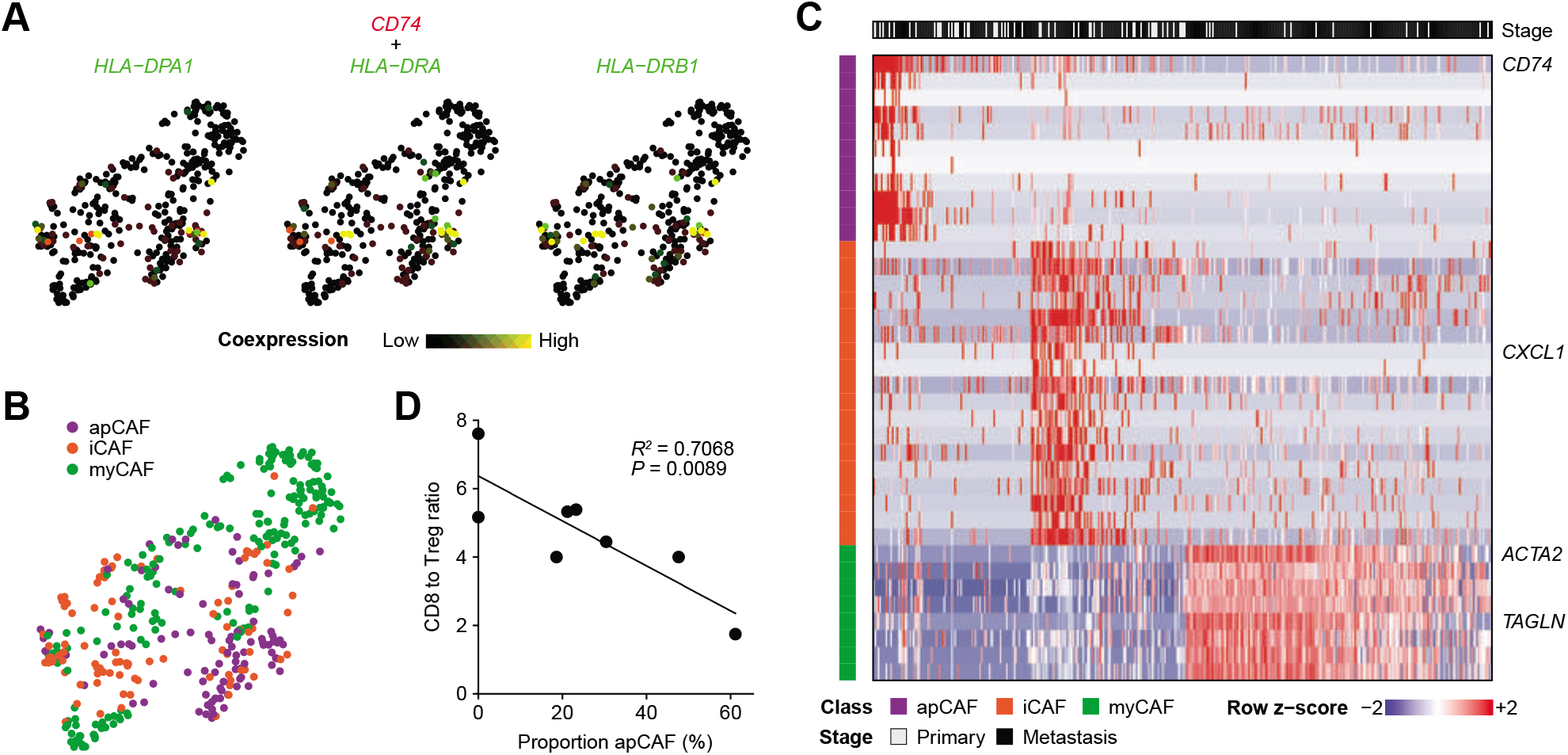
Stromal heterogeneity of PDAC is captured by biopsies. **A** Co-expression of selected apCAF marker genes projected onto UMAP plot of CAFs. *CD74* expression is represented by black (low) to red (high), whereas the expression of *HLA* genes is represented by black (low) to green (high). High expression of both *CD74* and variant MHC II molecule is shown as yellow. **B.** CAF subtypes from nearest template prediction projected onto UMAP plot. **C.** Heatmap showing scaled expression of apCAF, iCAF, and myCAF marker genes ordered by subtype. **D.** Correlation plot between apCAF abundance in each sample and the ratio of CD8 T cells (memory and effector) to regulatory T (Treg) cells.

#### PDAC-Infiltrating Immune Cells Acquire an Immune Suppressive Signature

PDAC is characterized by immunosuppressive microenvironment (1). To understand how immune cells change from the circulation to the PDAC TME, we performed scRNA-seq on peripheral blood mononuclear cells (PBMCs) from 8 of 9 patients obtained at or around the time of their tissue biopsies. Global comparison of all cell types revealed that there was a subpopulation of myeloid cells that were found only in the PDAC but not in the periphery (**Figure 5A; Supplementary Figure S5A**), and sub-clustering analysis revealed this unique population to be macrophages (**Figure 5B; Supplementary Figures S5B and S5C**). Given the potential polarization of tumor-associated macrophages into classical M1 and alternative M2 phenotypes (28), we performed trajectory inference analysis on monocytes and macrophages from PBMCs and tumors, which revealed a single unbranched trajectory representing the differentiation from monocytes to macrophages (**Figure 5C**). The expression of immune suppressive genes such as *MARCO* and *TREM2* increased with pseudotime (**Figure 5D**). Global gene expression changes along the pseudotime trajectory revealed upregulation of macrophage activation, acute and chronic inflammation, and negative regulation of myeloid-mediated immunity pathways (**Figure 5E**). Furthermore, macrophages were characterized by an upregulation of angiogenesis and hypoxia pathways (**Figure 5E**), which are associated with the M2 phenotype (28), confirming the immune suppressive role of tumor-infiltrating m acrophages.

**Figure 5.**
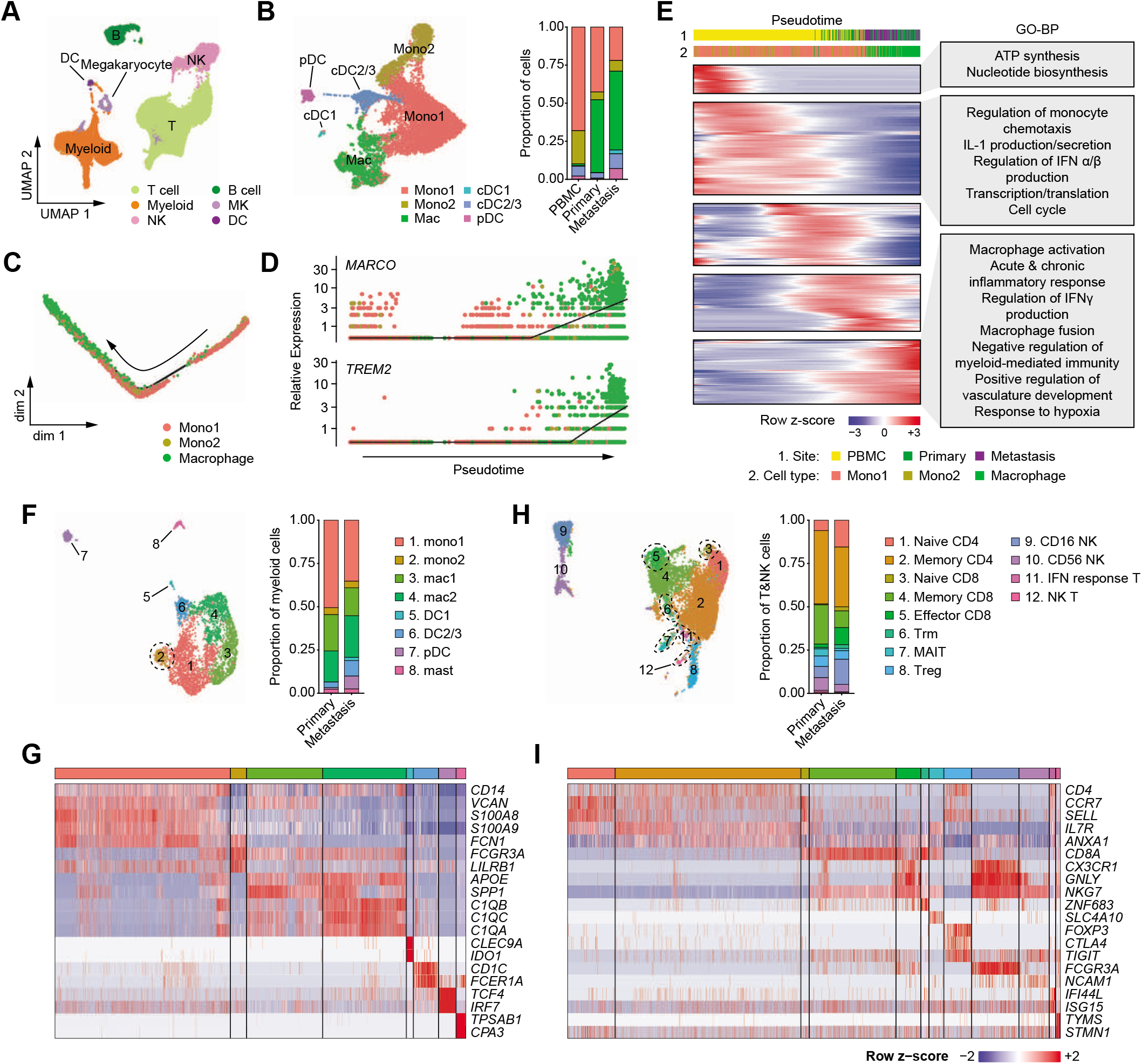
Acquisition of immune suppressive phenotype by tumor-infiltrating immune cells. **A.** UMAP plot of immune cells from peripheral blood mononuclear cells (PBMC) and tumors colored by cell type. **B.** UMAP of myeloid cells re-clustered from **A** colored by cell type (left) and bar plot showing proportions of each myeloid cell type as percent of all cells in PBMC, primary, and metastatic samples (right). **C.** Pseudotime trajectory of monocytes and macrophages. **D.** Dot plot representing the expression of *MARCO* (top) and *TREM2* (bottom) along the pseudotime trajectory from D colored by cell type. Solid black line indicates the mean expression at a given pseudotime. **E.** Heatmap showing dynamic gene expression changes through pseudotime (left) and enrichment of Gene Ontology biological processes (GO-BP) terms for selected gene cluster (right). **F.** UMAP of myeloid cells re-clustered from **Figure 1A** (left) and bar plot showing proportions of myeloid cell types in pooled primary and metastatic samples (right). Mono, monocyte; mac, macrophage; DC, dendritic cell; pDC, plasmacytoid DC. **G.** Heatmap showing scaled expression of select marker genes in each myeloid sub-cluster. **H.** UMAP of T and NK cell re-clustered from **Figure 1A** (left) and bar plot showing proportions of T and NK cell types in pooled primary and metastatic samples (right). Trm, tissue resident memory; MAIT, mucosal-associated invariant T; Treg, regulatory T; IFN, interferon. **I.** Heatmap showing scaled expression of select marker genes in each T and NK subcluster.

To elucidate the different types of myeloid cells present in the PDAC TME, we performed sub-clustering analysis of tumor-infiltrating myeloid cells and found 8 distinct cell types (**Figure 5F; Supplementary Table S6**). Using previously published subtyping of monocytes and dendritic cells (DCs) based on scRNA-seq profiles (29), we classified these sub-clusters as monocyte 1 (equivalent to CD14^high^ cells), monocyte 2 (equivalent to CD16^high^ cells), macrophage 1 (expressing *APOE* and *SPP1*), macrophage 2 (expressing complement molecules), DC1 (*CLEC9A*), DC2/3 (*CD1C*), plasmacytoid (p)DCs (*TCF4, IRF7*), and mast cells (*TPSAB1, CPA3*; **Figure 5G; Supplementary Figure S5D**). All sub-clusters were represented in primary tumors and metastases without significant differences in their proportions (**Supplementary Figure S5E**). In addition to *MARCO* and *TREM2*, tumor-associated macrophages expressed *PPARG* and *IL6* transcripts (**Supplementary Figure S5F**), the products of which are also implicated in immune suppression (30). Of note, a recent study comparing the immune compartments of PDAC and lung cancer revealed reduced numbers of DC infiltration in PDAC that led to the attenuation of anti-tumor T cell activity (31). Consistent with these findings, DCs represented smallest subsets of myeloid cells in our samples. Moreover, given that the DC1 subset expressed *IDO1* (**Figure 5G**), which inhibits T-cell proliferation (10,32), the DC1 subset may contribute to the immune suppressive phenotype in PDAC.

Analysis of tumor-infiltrating T and natural killer (NK) cells revealed 12 distinct sub-clusters (**Figure 5H; Supplementary Table S7**). Based on known markers for T and NK cells (33,34), we identified memory (*ANXA1*) and naïve (*SELL, CCR7*) CD4 T cells, memory and effector (*CX3CR1*) CD8 T cells, naïve CD8 T cells, mucosal-associated invariant T (MAIT) cells (*SLC4A10*), tissue resident memory CD8 T (Trm) cells (*ZNF683*), Tregs (*FOXP3, CTLA4*), CD56 NK cells, CD16 NK cells, IFN-response T cells (*IFI44L, ISG15*), and NK T cells (*TYMS, MKI67*; **Figure 5I; Supplementary Figure S6A**). Many T cells expressed inhibitory receptors *PDCD1, LAG3*, and *BTLA* (**Supplementary Figure S6B**), suggesting that T cells are present in the PDAC TME but their functions may be inhibited. Next, we compared T and NK cells in PBMCs and tumors (**Supplementary Figures S6C-S6E**). There was a trend of naïve CD4 T cells decreasing in proportion from periphery to primary to metastatic tumors, whereas memory CD4 and CD8 T cells showed a reverse trend (**Supplementary Figure S6F**), likely reflecting antigen exposure and creation of memory T cell populations predating metastatic spread. GSEA of tumorinfiltrating T and NK cells revealed the enrichment of hypoxia and negative regulation of immune response pathways (**Supplementary Figure S6G**), demonstrating the acquisition of immune suppressive phenotype similar to tumor-associated macrophages.

### Prediction of the PDAC Ligand-Receptor Interactome Reveals Multiple Immune Regulatory Pathways and Potentially Actionable Nodes

scRNA-seq has been used to predict potential ligand-receptor (LR) interactions, which we utilized to reveal potential mechanisms by which the crosstalk between tumor cells and immune cells may contribute to immunosuppression in PDAC. We applied a average expression-based LR interaction prediction (35,36) to assess potential relationships between major cell types of two primary (P3, P4) and two metastatic (liver, lung) tumors, as they contained the highest cell numbers and represented the most cellular diversity. We did not combine multiple samples for analysis to rule out potentially false interactions (e.g. T cell from liver metastasis to epithelial cells in a primary tumor). Global overview of the interactomes revealed that myeloid and dendritic cells were the most connected, and epithelial cells interacted closely with these two cell types (**Figures 6A and 6B; Supplementary Figures S7A and S7B**). One of the most significant LR pairs responsible for these connections was predicted to be between macrophage migration inhibitory factor (MIF) and the transcript for the invariant HLA-DR allele, CD74 (**Figure 6C**). This interaction has been implicated in promoting myeloid-derived suppressor cell-mediated immunosuppression in melanoma and breast cancer (37,38), while tumor cell MIF expression has been linked to a more aggressive phenotype in PDAC (39). Another prevalent interaction between tumor epithelial cells and myeloid/dendritic cells involved amyloid precursor protein (APP) and CD74; of interest, we have previously described that APP overexpression in PDAC cells upregulates proliferation (40). The potential role of neoplastic APP in subverting antigen presentation via CD74 interaction on immune cells in the PDAC TME is uncharted.

**Figure 6.**
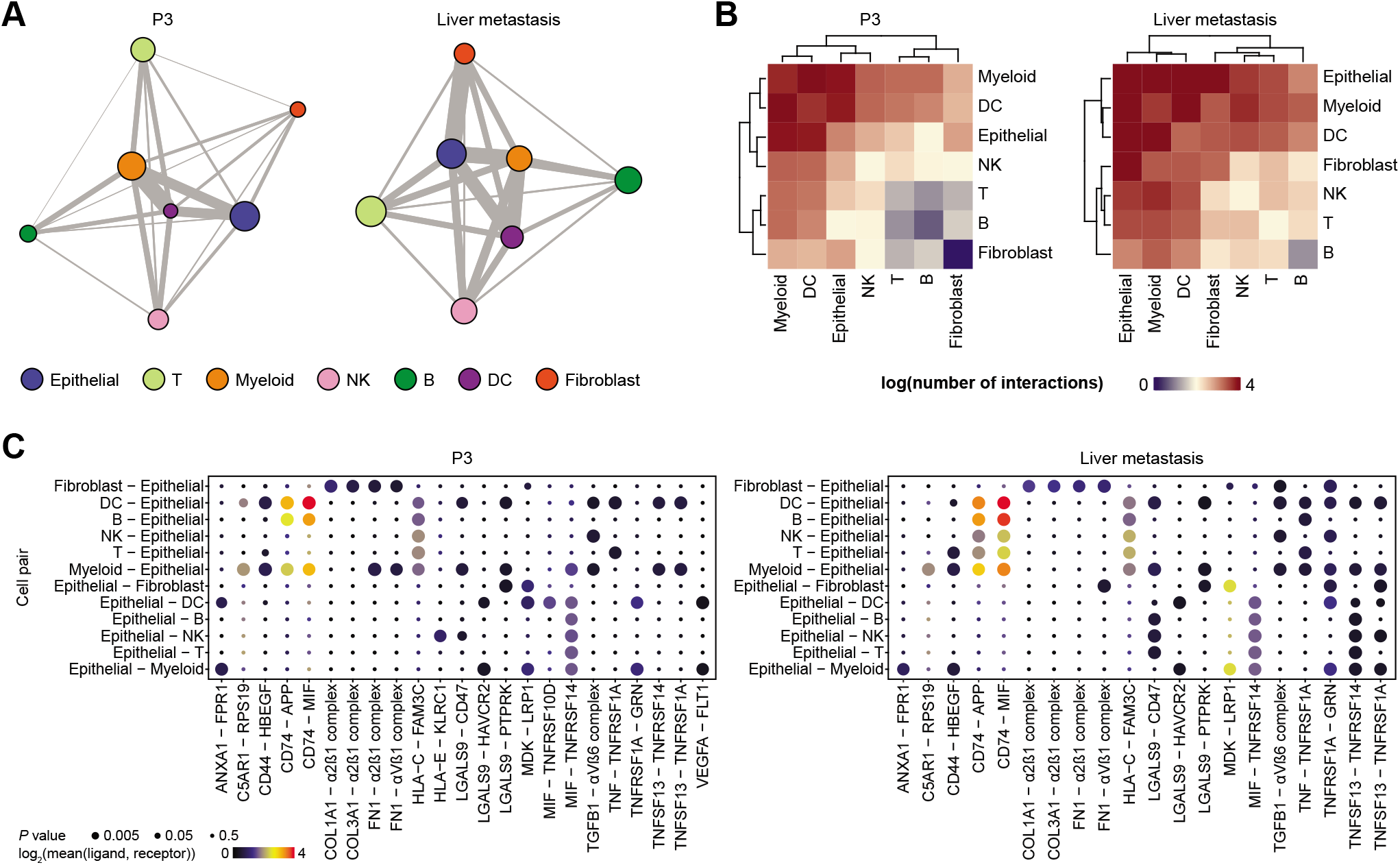
Ligand-receptor interaction predictions between major cell types. **A.** Network plots of a primary PDAC (P3, left) and liver metastasis (right) demonstrating potential ligand-receptor interactions. Each node represents a cell type and size reflects relative number of cells. Each edge represents the number of significant interactions between each cell-type pair and its thickness is proportional to the number of interactions. **B.** Heatmaps of log-transformed number of significant interactions between cell-type pairs in a primary PDAC (P3, left) and liver metastasis (right). **C.** Bubble plots representing top significant ligand-receptor interactions between different cell-type pairs involving epithelial cells in a primary PDAC (P3, left) and liver metastasis (right). Size of each bubble represents *P* value and color represents the mean expression of ligand and receptor genes.

Given that scRNA-seq data can be biased by gene dropout, thereby diminishing the average expression of genes in each cell type, we next predicted LR interactions at single-cell level (41). When looking at predicted interactions involving ligands from cells in the TME and cognate receptors on epithelial cells, we found that epithelial cells were more strongly connected to fibroblasts and myeloid cells, whereas there were fewer connections with T and B cells (**Figure 7A; Supplementary Figures S7C and S7D**). Many predicted interactions between epithelial cells and fibroblasts involved components of the ECM (**Figure 7B**), which is expected given that one of the major functions of fibroblasts is the production of ECM proteins. There were many pro-inflammatory interactions such as IFNG from T/NK cells and IFNGR1/2 on epithelial cells (**Figure 7B**), once again pointing to inflammatory signals present in the PDAC microenvironment.

**Figure 7.**
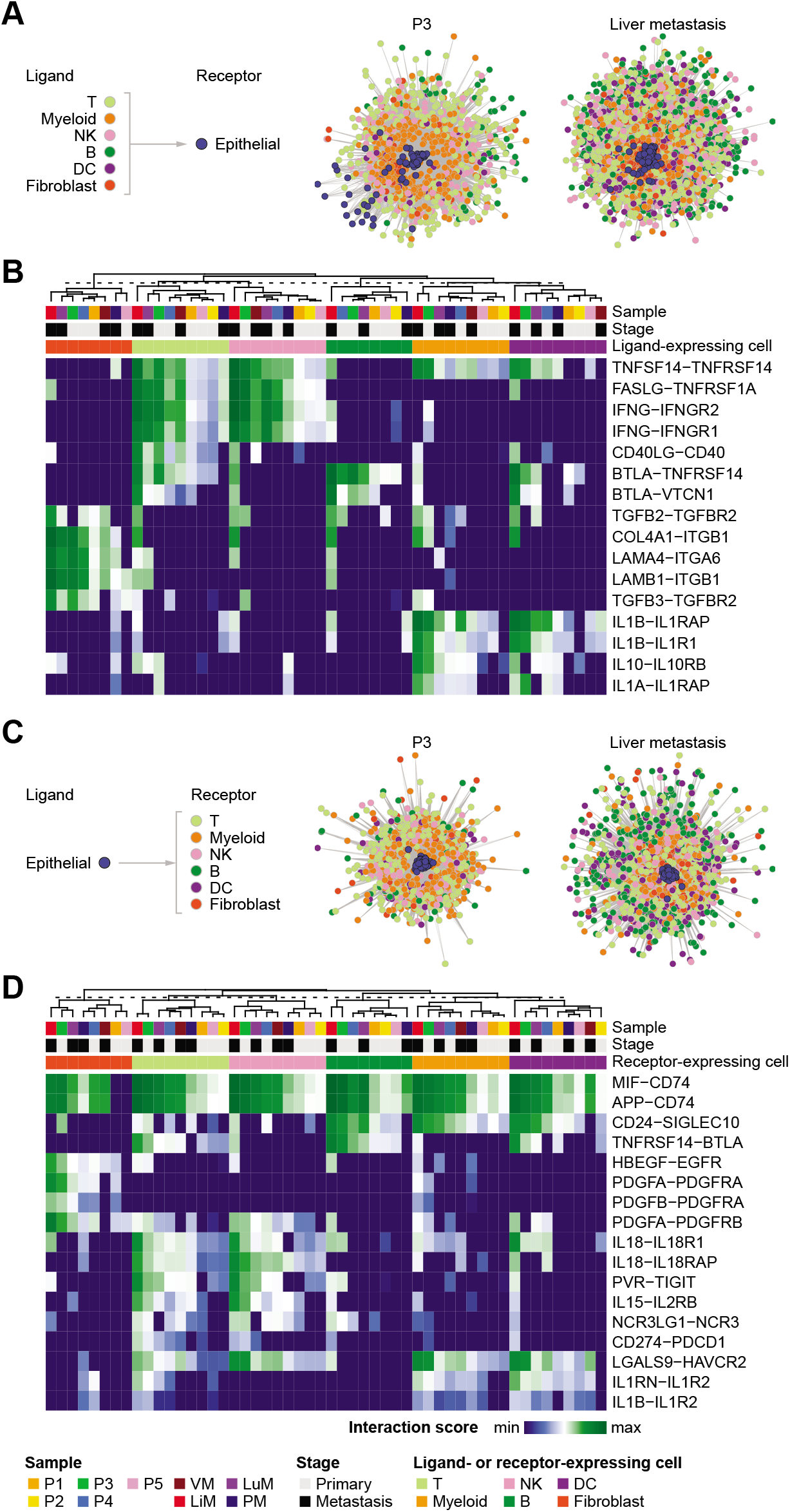
Ligand-receptor interaction predictions between single cells. **A.** Network plots of a primary PDAC (P3, left) and liver metastasis (right) demonstrating potential ligand-receptor interactions in which epithelial cells express the receptor. Each node represents a single cell and the edge represents the number of ligand-receptor pairs between two cells. **B.** Heatmap showing top interaction scores (normalized number of ligand-receptor interactions) for each cell-type pair where epithelial cells express the receptor. Sample origin, stage, and ligand-expressing cells are annotated above the heatmap. See bottom of figure for legend. **C.** Network plots of a primary PDAC (P3, left) and liver metastasis (right) demonstrating potential ligand-receptor interactions in which epithelial cells express the ligand. **D.** Heatmap showing top interaction scores for each cell-type pair where epithelial cells express the ligand. Sample origin, stage, and receptor-expressing cells are annotated above the heatmap. Ligand-receptor interaction class is annotated to the left. P1-5, primary 1-5; VM, vaginal apex metastasis; LiM, liver metastasis; LuM, lung metastasis; PM, peritoneal metastasis.

We also explored the reverse interactions in which ligands were expressed on epithelial cells and receptors on TME cells, which again revealed more connections with fibroblasts and myeloid cells (**Figure 7C; Supplementary Figure S7E and S7F**). Most prominent interactions with fibroblasts were growth factors, and interactions with immune cells involved cytokine and checkpoint genes (**Figure 7D**). Of note, epithelial cells interacted with T/NK cells via IL18-IL18R1 and LGALS9-HAVCR2. IL18 has been shown to promote IFNγ production in T/NK cells (42), and the interaction between LGALS9 and HAVCR2 can suppress T cell-mediated immunity (43). Additional immunosuppressive interactions between epithelial and T cells included PVR-TIGIT and CD274-PDCD1. Interactions with myeloid and dendritic cells included CD24-SIGLEC10, which promotes immune evasion and is a potential target in cancer immunotherapy (44). One of the most common interactions between B and epithelial cells was BTLA-TNFRSF14 (also known as herpesvirus entry mediator or HVEM), which can suppress both T and B cell-mediated immunity (45). Overall, these results suggest that PDAC tumor cells can drive an immunosuppressive microenvironment by affecting multiple immune cell types. MIF-CD74 and APP-CD74 interactions were abundant between epithelial cells and multiple TME cell types, denoting high expression of MIF and APP in PDAC tumor cells and CD74 in TME cells with a strong potential for *in vivo* functionality.

## Discussion

PDOs are a promising tool in precision medicine approaches for PDAC, but molecular alterations associated with prolonged *ex vivo* culture conditions remain less well characterized. We present evidence of transcriptomic evolution of PDOs during *ex vivo* establishment and passaging. The differences were more pronounced with the transition from tumor tissue to PDO, which was characterized by the loss of inflammatory microenvironmental cues such as TNF and IFN signaling and gain of proliferative pathways. This transcriptomic shift led to altered drug sensitivity predictions on scRNA-seq data, although there was an “invariant” set of compounds that were predicted in common with the contemporaneous biopsy sample, which might reflect therapies more likely to be effective *in vivo*.

Given these potential limitations of organoids, it is imperative to develop orthogonal methods to comprehensively characterize limited biopsy specimens in clinically informative ways. To that end, we systematically analyzed scRNA-seq from “real world” biopsies of primary and metastatic PDAC, demonstrating that even these scant samples recapitulate significant cellular heterogeneity. Our data showed higher proportion of epithelial cells compared to a recent scRNA-seq studies on surgically resected PDAC tumors (10,11), while the most common immune cell types were myeloid and T cells, consistent with these studies. We performed transcriptomics subtyping at single-cell resolution, which revealed two subtypes specific to tumor epithelial cells, namely classical (or pancreatic progenitor) and basal-like (or QM/squamous), confirming the results from a previous analysis of bulk tumors (20). Metastatic lesions were more likely to harbor a higher proportion of aggressive basal-like subtype of cells, which was also previously reported in a separate PDAC cohort (46). Notably, we show that although bulk RNA sequencing classifies PDAC dichotomously as one of two subtypes, at single cell resolution, even limited PDAC biopsies are comprised of more than one subtype. This finding may be of clinical importance in light of the COMPASS trial, which showed that classical PDAC is more likely to respond to first-line FOLFIRINOX, while basal-like tumors tend to be chemo-resistant (47). The existence of basal-like cells within an “apparently classical” tumor (as deciphered by bulk sequencing) could lead to preferential enrichment of basal-like cells and rapid emergence of resistance. Thus, future iterations of trials like COMPASS that rely on subtyping using core biopsies might need to provide a quantitative measure of cellular subtypes present, rather than a qualitative dichotomous readout.

Furthermore, the overall gene expression profile and cell type proportions revealed several putative mechanisms of immune suppression in PDAC, such as the expression of immune suppressive genes by myeloid cells and inhibitory receptors in T cells, polarization of macrophages towards M2 phenotype, as well as the presence of apCAFs that may inhibit T cell-mediated immune response. Prediction of the prevalent ligand-receptor “interactome” also revealed multiple mechanisms by which tumor-TME interactions may contribute to inflammation and immune evasion.

Through integrative analysis of scRNA-seq data, we uncovered implicit heterogeneity of low-input biopsy samples in PDAC that demonstrates potential mechanisms for immune evasion and provide high-resolution data correlating cell types to clinically relevant molecular subtypes. Given the potential limitations of organoids in maintaining the original tumor transcriptomic program and predicting response to therapy across passages, deep characterization using a contemporaneous biopsy may provide a valid orthogonal approach to precision medicine approaches.

## Methods

### Sample Acquisition

A total of 9 patients were recruited at the University of Texas MD Anderson Cancer Center through informed written consent following institutional review board approval (Lab00-396 and PA15-0014). The study was conducted in accordance with Good Clinical Practices concerning medical research in humans per the Declaration of Helsinki. Five primary pancreatic cancers and four metastatic lesions in liver, lung, peritoneum, and vaginal apex were used. Histologic confirmation of PDAC was performed by a pathologist.

### Single Cell Dissociation

Pancreatic cancer biopsies were collected in high-glucose DMEM supplemented with GlutaMAX, HEPES buffer, and 1% bovine serum albumin (BSA; all Thermo Fisher). Samples were delivered to the laboratory within 2 hours after the procedure. Single cell dissociation was performed as previously described (9). Briefly, samples were minced with sterile surgical scalpels to 0.5-1 mm fragments in approximately 1 ml of media. After centrifugation for 5 minutes at 125 g, cells were resuspended in DMEM with 0.5 mg/ml Liberase TH (Sigma-Aldrich) and 1% penicillin-streptomycin (Corning) and incubated in an orbital shaker for 15 min at 37°C. Liberase was quenched with equal volume of 1% BSA in DMEM followed by centrifugation for 5 minutes at 125 g. Cells were resuspended in 2 ml Accutase (Sigma-Aldrich) and incubated in an orbital shaker for 15 minutes at 37°C. Organoids were dissociated by incubating with TrypLE Express (Thermo Fisher) for 5-15 minutes at 37°C followed by manual disruption. Dissociated cells were passed through a 40 μm strainer and centrifuged for 5 minutes at 125 g. Isolated cells were resuspended in 0.04% BSA in phosphate buffered saline (Corning) for subsequent viability analysis and counting.

### Single-Cell RNA Library Preparation and Sequencing

Single-cell transcriptomic amplification and library preparation was performed using the 5’ or 3’ Library Construction Kit (10x Genomics) following the manufacturer’s recommendations. Final library quality and concentration were measured on TapeStation System using High Sensitivity D5000 ScreenTape (Agilent). Libraries were sequenced on NextSeq 500 (Illumina) according to the manufacturer’s instructions.

### Sequencing Data Processing and Analysis

Raw base call (BCL) files were demultiplexed and converted to FASTQ files, which were subsequently used to generate feature-barcode matrices using Cell Ranger RNA v3.1 (10x Genomics) and hg19 was used as a reference. Additional analytical methods are included in the Supplementary Information.

## Acknowledgements

We thank Senthil K. Muthuswamy (Beth Israel Deaconess Medical Center) for providing the progenitor and tumor organoid media (PaTOM) and Mark W. Hurd (The University of Texas MD Anderson Cancer Center) for assistance with sample acquisition.

