## Supplementary File for "Elucidation of tumor-stromal heterogeneity and the ligand-receptor interactome by single cell transcriptomics in real-world pancreatic cancer biopsies"

#### Supplementary Methods

##### Organoids Generation

Dissociated cells (as described in Methods) were resuspended in progenitor and tumor organoid media (PaTOM) (1) and centrifuged for 5 min at 125 g. Cell pellet was resuspended in Growth Factor-Reduced Matrigel (Corning) and plated as a dome in a Nunclon Delta surface treated 24-well plate (Thermo Fisher). Cells embedded in Matrigel were incubated for 15 min at 37°C to allow polymerization and covered with PaTOM. Organoids were maintained in a 37°C incubator with 5% $CO_2$.

##### Peripheral Blood Mononuclear Cell (PBMC) Isolation

Whole blood samples were collected in acid citrate dextrose (ACD) tubes (BD Biosciences) and processed within 4 hours of phlebotomy. To isolate PBMCs, whole blood was centrifuged for 10 min at 2,500 g, plasma was removed, and the remaining fraction was layered on Lymphocyte Separation Media (Corning) followed by centrifugation for 30 min at 620 g. PBMCs were stored at -80°C in CryoStor CS10 (STEMCELL Technologies) until ready for use.

**Single-Cell Copy Number Sequencing and Analysis**

Tissues and organoids were dissociated into single cell suspension in 0.04% BSA/PBS as described in Methods. Single cell copy number libraries were generated and sequenced using Single Cell CNV Kit (10x Genomics) following the manufacturer's recommendations. All CNV libraries were sequenced using NextSeq 500 (Illumina) using 300-cycle, paired-end, single-indexing protocol (read 1 = 150, read 2 = 150, i7 index = 8 cycles, i5 index = 0 cycles). Raw sequencing reads were aligned to hg19 reference genome and processed using Cell Ranger DNA v1.1 (10x Genomics).

**Dimensionality Reduction and Cell Clustering**

Initial single-cell analysis was performed using Seurat v3.1.0 (2). Cells containing less than 200 genes were removed, and log-normalized expression of 2,000 variable genes were used to reduce the data into two-dimensional space. Cells from all 9 samples were combined using FindIntegrationAnchors and IntegrateData functions. Cell identities were manually assigned using highly expressed genes. To retain the highest number of cells for subsequent analyses, no additional filters were initially applied. To analyze the heterogeneity within each cell type, cells from each cluster were subsetted and reanalyzed using 2,000 variable genes while keeping the integration. Cells with higher expression of mitochondrially encoded genes were filtered out *post hoc* based on the results of the sub-clustering.

**Cell State Trajectory Analysis**

Trajectory inference was performed using Monocle v2.13.0 (3). Analysis was done using raw counts obtained from Seurat objects with preserved cellular metadata. Top 1,000 variable genes were used to reduce the data and order the cells. Pseudotime heatmap was created using genes with dynamic expression changes obtained using differentialGeneTest function specifying fullModelFormulaStr = "~sm.ns(Pseudotime)" and q-value cutoff of $1 \times 10^{-10}$ for myeloid cells and $1 \times 10^{-40}$ for epithelial cells. Branched pseudotime heatmap was created using BEAM function and q-value cutoff of $1 \times 10^{-5}$. Genes were clustered based on pattern of expression changes and extracted from Monocle object for subsequent analysis.

**Gene Expression Classifier**

Molecular subtyping was performed with nearest template prediction algorithm and cosine correlation. CMScaller R package v0.99.1 (4) was used as a wrapper to implement nearest template prediction using log-normalized counts from Seurat object and previously published gene lists (5-7). False discovery rate (FDR) of 0.2 was used as a cutoff to determine the significance of classification (4). Pseudo-bulk RNA expression was projected by taking the summation of raw counts across all genes for each sample prior to applying any filters. The score for each subtype represents mean expression of genes belonging to that subtype.

**Gene Set Enrichment Analysis (GSEA)**

To obtain significantly enriched pathways, marker genes were first obtained using FindMarkers function in Seurat. Resulting genes were ranked by -log10(FDR) multiplied by fold-change directions so that upregulated genes had positive values and downregulated genes had negative values. Pre-ranked GSEA was performed using fgsea R package v1.12.0 (8) specifying 10,000 permutations. Gene ontology (GO) enrichment analysis was done on geneontology.org (9,10), and only positively enriched terms with FDR < 0.05 were included.

**Drug Sensitivity Prediction and Validation**

The next generation connectivity map was used to predict drug sensitivity from scRNA-seq expression (11). Briefly, marker genes for individual subclusters or the pseudo-bulk profile for tissue and organoids were obtained from Seurat. Human pancreatic ductal cells from a previous droplet-based scRNA-seq data on normal pancreas (12) were used as reference to measure differentially expressed genes. Genes were ranked by average log-fold change and filtered by adjusted P value (< 0.05). A list of up to 150 upregulated and downregulated genes (total of up to 300 genes) were used as input for the connectivity map (clue.io). Connectivity scores greater than 90 or less than -90 were used as cutoffs to predict resistance and sensitivity, respectively. To validate the sensitivity prediction to gemcitabine, organoids were dissociated into single cells as described in Methods and plated in untreated round bottom 384-well plates with opaque walls (Corning) at a density of 1,000 cells/well in 50 μl in PaTOM media (day 0) and maintained at 37°C with 5% $CO_2$. Cells were treated with gemcitabine (Selleck) at noted concentrations from days 3 to 5, changing the media daily. On day 6, cell viability was assessed using CellTiter-Glo 3D Cell Viability Assay (Promega) per the manufacturer's protocol and luminescence was read on Cytation 3 Cell Imaging Multi-Mode Reader (BioTek).

**Copy number inference from scRNA-seq**

Copy number alterations from scRNA-seq were inferred using InferCNV v1.0.4 (https://github.com/broadinstitute/inferCNV) and cutoff was set to 0.1. Non-epithelial cells were used as reference.

**Ligand-Receptor Interaction Prediction**

CellPhone DB v2.1.1 (13,14) was used to analyze cell-type based ligand-receptor interactions. Log-normalized counts from filtered Seurat objects and major cell type metadata (epithelial, T, myeloid, NK, etc.) were used as input. Method statistical_analysis and --threshold=0.25 were used. Interaction network analysis between individual cells was modeled after a previous study (15) using a previously curated ligand-receptor pairs (16) with the addition of MIF-CD74, APP-CD74, and PVR-TIGIT. Only ligand or receptor genes with Seurat log-normalized expression value greater than 0.5 were used. Up to 100 cells were randomly subsetted from each cell type without setting a seed, and 5 iterations were performed to test reproducibility. Only those ligand-receptor interactions contributing to at least 25 cell pairs were counted. We summed the number of ligand-receptor pairs for each cell-cell interaction and normalized the interactions by dividing by the maximum number of interactions in all cell pairs. The igraph R package (v1.2.4.1) was used to create network plots applying an edge threshold (minimum) of 0.1 and nodes with at least 3 edges. Layout was set to Fruchterman-Reingold with weight specified as relative number of interactions between each cell pair.

Network heatmap was made by counting the number of ligand-receptor pairs for each cell-type pair. The interaction score was calculated by normalizing the number of ligand-receptor interactions for each pair of cell types with the total number of cells involved in the cell pairs followed by log transformation.

**Supplementary Figure Legends**

**Supplementary Figure S1. Evolution of patient-derived organoids (PDOs).**

**A.** UMAP plot of cells from PDO-1 at passage 5 (green) and passage 15 (purple, navy).

**B.** Pseudotime trajectory of single cells from passage 5 (green) and passage 15 (purple, navy).

**C.** Branched heatmap showing dynamic gene expression (left). Pseudotime progresses from left to right. Enriched Gene Ontology biological process (GO-BP) terms for each gene cluster are listed on the right.

**D.** Heatmap showing inferred copy number alterations (CNA) of early and late passage PDO-1. Cells (rows) are ordered by pseudotime. Red represents copy number gains and blue represents copy number losses.

**E**. Heatmap showing single cell copy numbers from vaginal metastasis tissue (PDAC-VM, top) and PDO-2 (bottom).

**F.** Heatmap showing inferred CNA of non-epithelial cells from PDAC-VM (top), epithelial cells from PDAC-VM (middle), and cells from PDO-2 (bottom). Red represents copy number gains and blue represents copy number losses.

**G.** UMAP plot of epithelial cells from PDAC-VM showing clusters used as input for drug sensitivity prediction.

**H.** UMAP plot of cells from PDO-2 showing clusters used as input for drug sensitivity prediction.

**I.** Venn diagram showing overlap of compounds predicted to target epithelial cells in

PDAC-VM (green) and PDO-2 (purple).

**Supplementary Figure S2. PDAC heterogeneity and epithelial heterogeneity from scRNA-seq of biopsies.**

**A.** Dot plot showing average proportions of each cell type as percent of all cells. Error bars represent S.E.M.

**B.** Heatmap showing mean expression of marker genes in each epithelial sub-cluster (left) and enrichment of GO-BP terms for each cluster of genes (right).

**C.** Dot plot showing average proportions of each epithelial cell sub-cluster as percent of all epithelial cells. Error bars represent S.E.M.

**D.** Heatmap showing inferred CNA of all cells from PDAC biopsies. Stromal cells are on top and epithelial cells are below. Cells were grouped by cell type then by sample of origin. Red represents copy number gains and blue represents copy number losses.

**Supplementary Figure S3. Molecular subtyping of single cells.**

**A-C.** Heatmaps of scaled expression of genes in Bailey (**A**), Collisson (**B**), and Moffit (**C**) subtypes for all cells (left) and epithelial cells (right). Rows represent genes and columns represent cells. Genes are ordered by subtype, and cells are ordered by subtype prediction followed by *P* value. Annotations above each heatmap show cell type, whether the cell is from a metastatic lesion, *P* values of subtype prediction in white bars, and scores for each subtype based on the mean expression of genes belonging to the corresponding subtypes.

**D.** Bar plots showing proportions of all cells (left) and epithelial cells (right) that classified into each classifier with false discovery rate (FDR) < 0.2.

**E.** Sankey diagrams showing relationships between cell types (from Figure 2A) on the left and molecular subtypes on the right. Only cells with subtype prediction FDR < 0.2 are included. The number of cells that were confidently assigned a subtype are displayed above each diagram.

**F.** Sankey diagrams showing relationships between epithelial cell types (from Figure 2C) on the left and molecular subtypes on the right. Only cells with subtype prediction FDR < 0.2 are included. The number of cells that were confidently assigned a subtype are displayed above each diagram.

**G.** Bar plot showing percent of epithelial cells in each sample (left) and heatmap representing molecular subtype prediction of projected pseudo-bulk RNA expression data (right).

**H.** Bar plots showing proportions of cells classifying into Bailey subtype with FDR < 0.2 in pooled primary and metastatic samples (left) and individual samples (right).

**I.** Bar plots showing proportions of cells classifying into Collisson subtype with FDR < 0.2 in pooled primary and metastatic samples (left) and individual samples (right).

**J.** Bar plots showing proportions of cells classifying into Moffitt subtype with FDR < 0.2 in pooled primary and metastatic samples (left) and individual samples (right).

**Supplementary Figure S4. Heterogeneity of cancer-associated fibroblasts (CAFs).**

**A.** UMAP plot of fibroblasts re-clustered from Figure 2A (top) and bar plot showing proportions of fibroblast subtypes in pooled primary and metastatic samples (bottom).

**B.** Violin plots representing relative expression levels of select CAF marker genes in the two initial CAF subtypes.

**C.** Dot plot showing average proportions of each fibroblast subtype as percent of all fibroblasts. Error bars represent S.E.M.

**D.** Violin plots showing relative expression of non-CAF marker genes in three CAF subtypes.

**Supplementary Figure S5. Heterogeneity of myeloid cells.**

**A.** UMAP plot of immune cells from PBMC and tumors colored by site.

**B.** UMAP of myeloid cells re-clustered from Figure 5A colored by site.

**C.** Bubble plot showing highly expressed marker genes in each myeloid cell type, with cell types in rows and genes in columns. Size of each bubble represents percent of cells expressing marker and color represents the level of expression.

**D.** Heatmap showing scaled expression of top marker genes for each myeloid sub-cluster.

**E.** Dot plot showing average proportions of each myeloid cell type as percent of all myeloid cells in primary and metastatic samples. *P* values are calculated by two-sided Mann-Whitney *U*-test. Error bars represent S.E.M.

**F.** Expression of select immune suppressive genes projected onto UMAP plot from Figure 5F (excluding pDC and mast cell sub-clusters).

**Supplementary Figure S6. Heterogeneity of T and NK cells.**

**A.** Heatmap showing scaled expression of top marker genes for each T and NK sub-cluster.

**B.** Expression of select inhibitory receptor genes projected onto UMAP plot from Figure 5H (excluding NK sub-clusters).

**C.** UMAP of T and NK cells re-clustered from Figure 5A colored by cell type.

**D.** UMAP of T and NK cells re-clustered from Figure 5A colored by site.

**E.** Bubble plot showing highly expressed marker genes in each T and NK cell type, with genes in rows and cell types in columns. Size of each bubble represents percent of cells expressing marker and color represents the level of expression.

**F.** Bar plot showing proportions of selected T cell types as percent of T and NK cells in PBMC, primary, and metastatic samples. Error bars represent S.E.M.

**G.** Bubble plot showing enrichment of pathways in tumor-infiltrating T and NK cells compared to peripheral cells, with cell types in columns and pathways in rows. Size of each bubble represents FDR and color represents NES.

**Supplementary Figure S7. Ligand-receptor interaction prediction in PDAC.**

**A.** Network plots of a primary PDAC (P4, left) and lung metastasis (right) demonstrating ligand-receptor interactions. Each node represents a cell type and size is reflective of number of cells. Each edge represents number of significant interactions between each cell type and thickness is reflective of number of interactions.

**B.** Heatmaps of a primary PDAC (P4, left) and lung metastasis (right) demonstrating the log-transformed number of significant interactions between each cell type pairs.

**C.** Network plots of a primary PDAC (P4, left) and lung metastasis (right) demonstrating potential ligand-receptor interactions with epithelial cells expressing the receptor. Each node represents a single cell and the edge represents the number of ligand-receptor pairs between two cells.

**D.** Heatmap showing relative number of interactions (normalized by maximum number of interactions) between epithelial cells and stromal cells where epithelial cells express the receptor. Each row represents epithelial cell from specified sample.

**E.** Network plots of a primary PDAC (P4, left) and lung metastasis (right) demonstrating potential ligand-receptor interactions with epithelial cells expressing the ligand. Each node represents a single cell and the edge represents the number of ligand-receptor pairs between two cells.

**F.** Heatmap showing relative number of interactions (normalized by maximum number of interactions) between epithelial cells and stromal cells where epithelial cells express the ligand. Each row represents epithelial cells from specified sample.

**Supplementary Table Legends**

**Supplementary Table 1.** Total cell counts summarized by cell type and sample type (primary or metastasis).

**Supplementary Table 2.** Summary of epithelial sub-cluster cell counts by sample type.

**Supplementary Table 3.** Results of molecular subtyping of all cells by Bailey, Collisson, and Moffitt classifiers. Numbers are organized by sample of origin and sample type.

**Supplementary Table 4.** Results of molecular subtyping of epithelial cells by Bailey, Collisson, and Moffitt classifiers. Numbers are organized by sample of origin and sample type.

**Supplementary Table 5.** Summary of fibroblast subtypes by sample type.

**Supplementary Table 6.** Summary of myeloid cell subtypes by sample type.

**Supplementary Table 7.** Summary of T and NK cell subtypes by sample type.

### Supplementary Figure S1

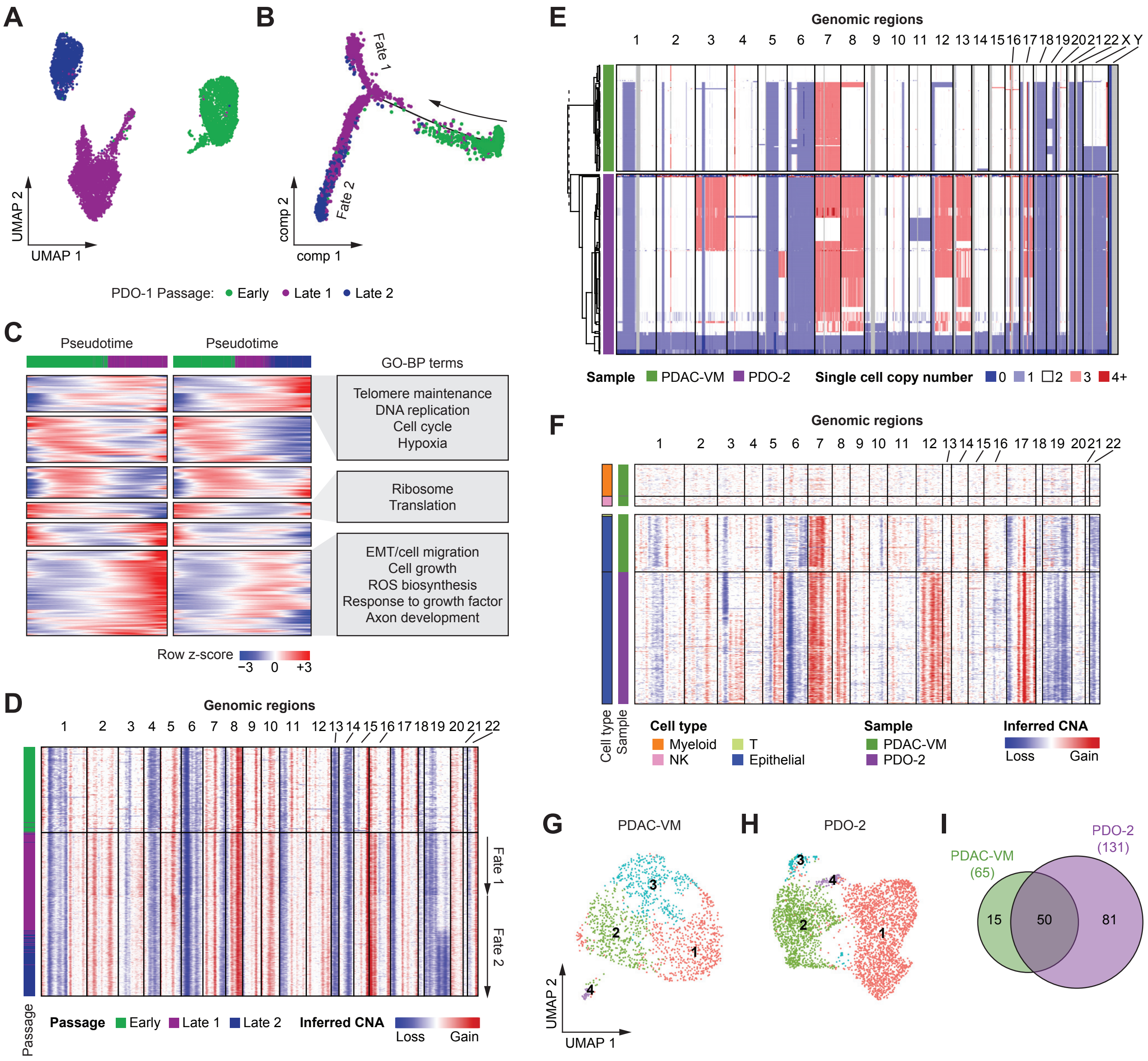

### Supplementary Figure S2

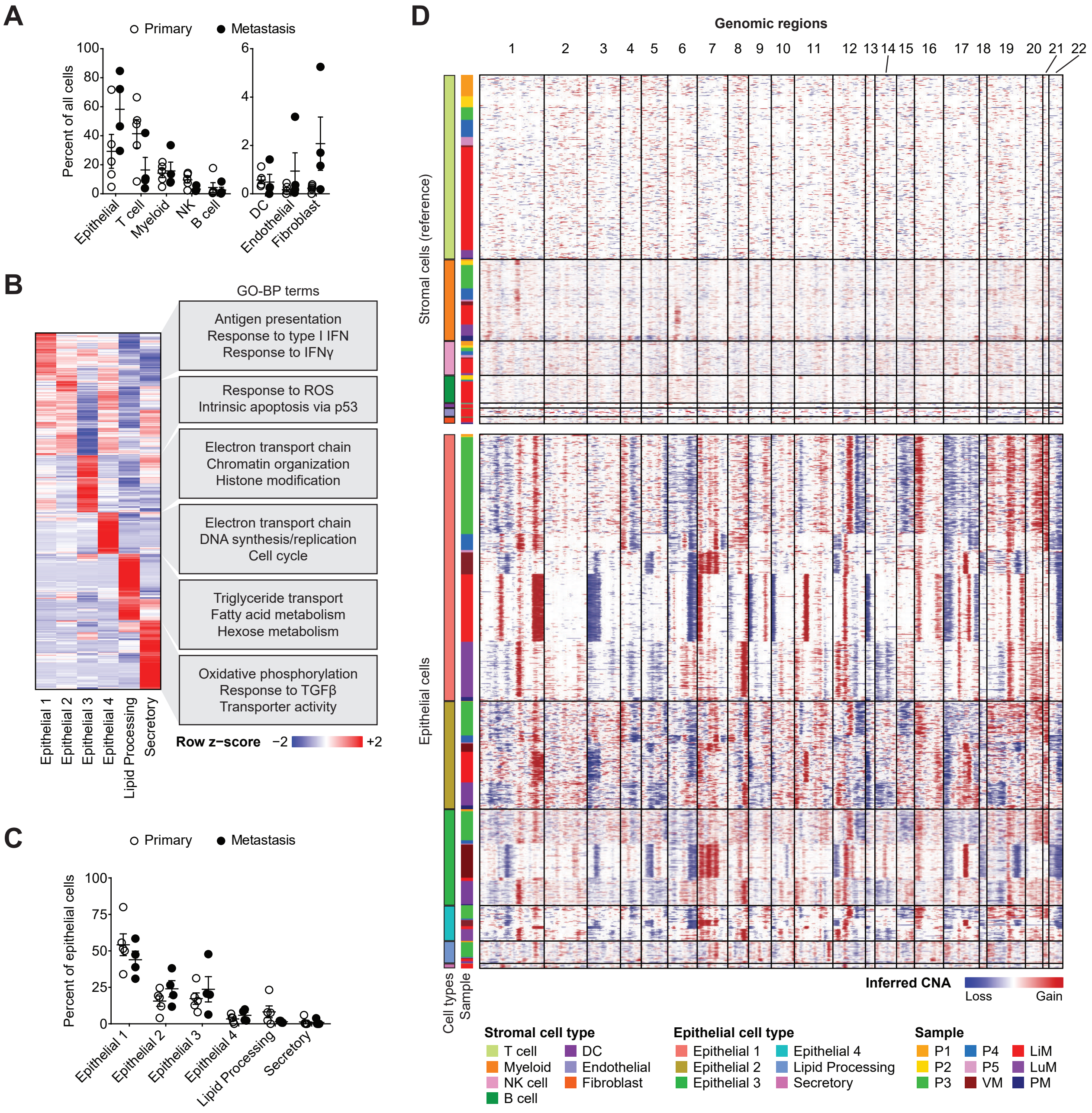

### Supplementary Figure S3

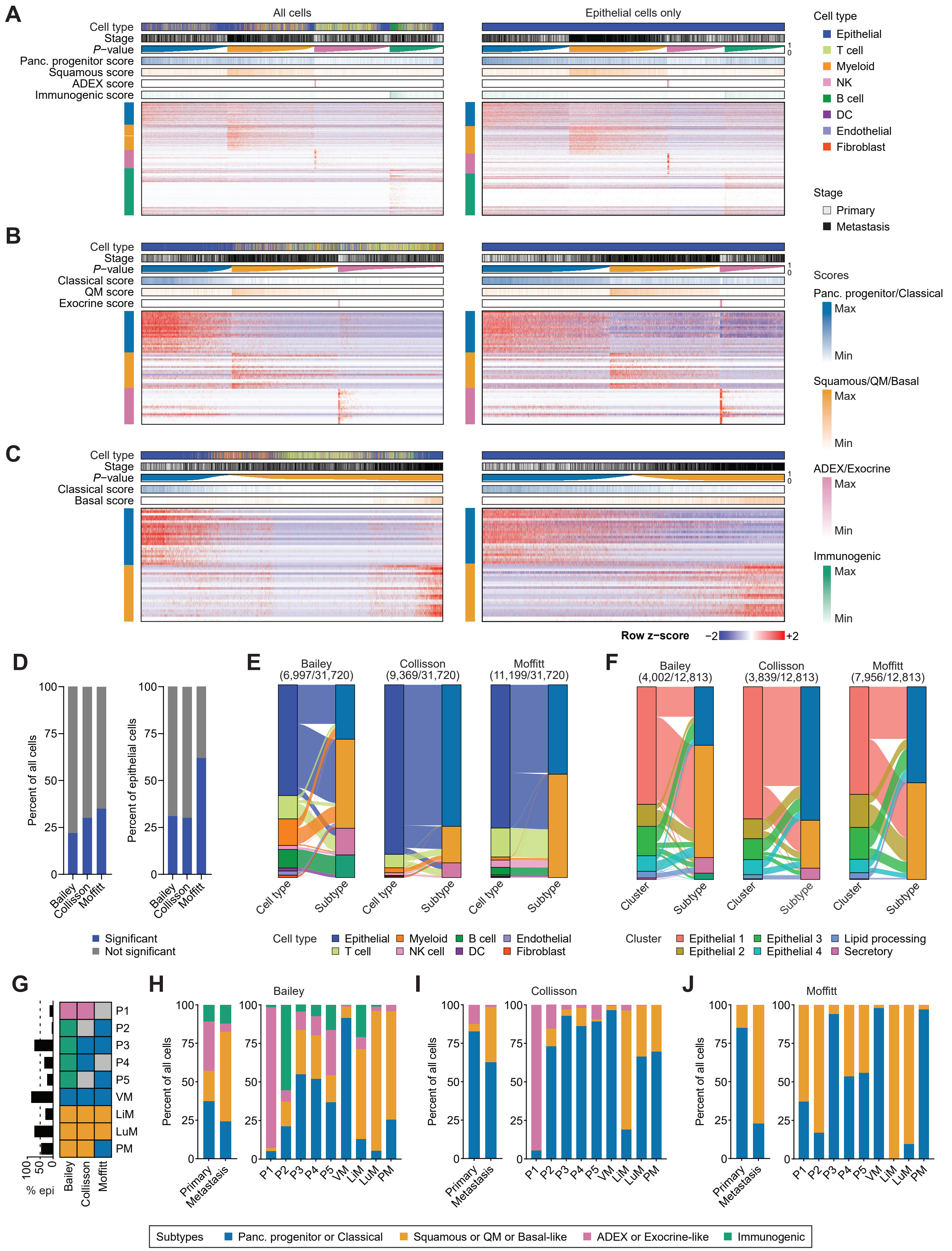

### Supplementary Figure S4

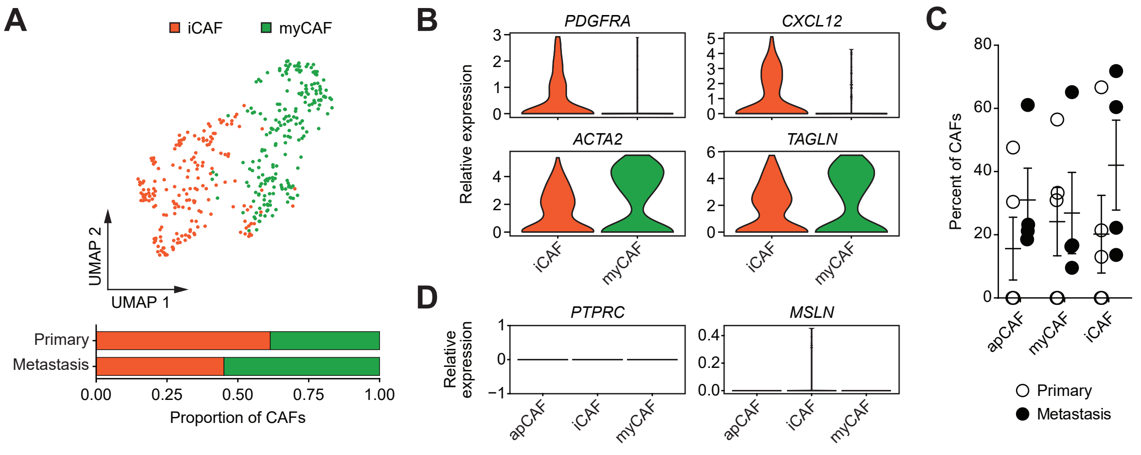

### Supplementary Figure S5

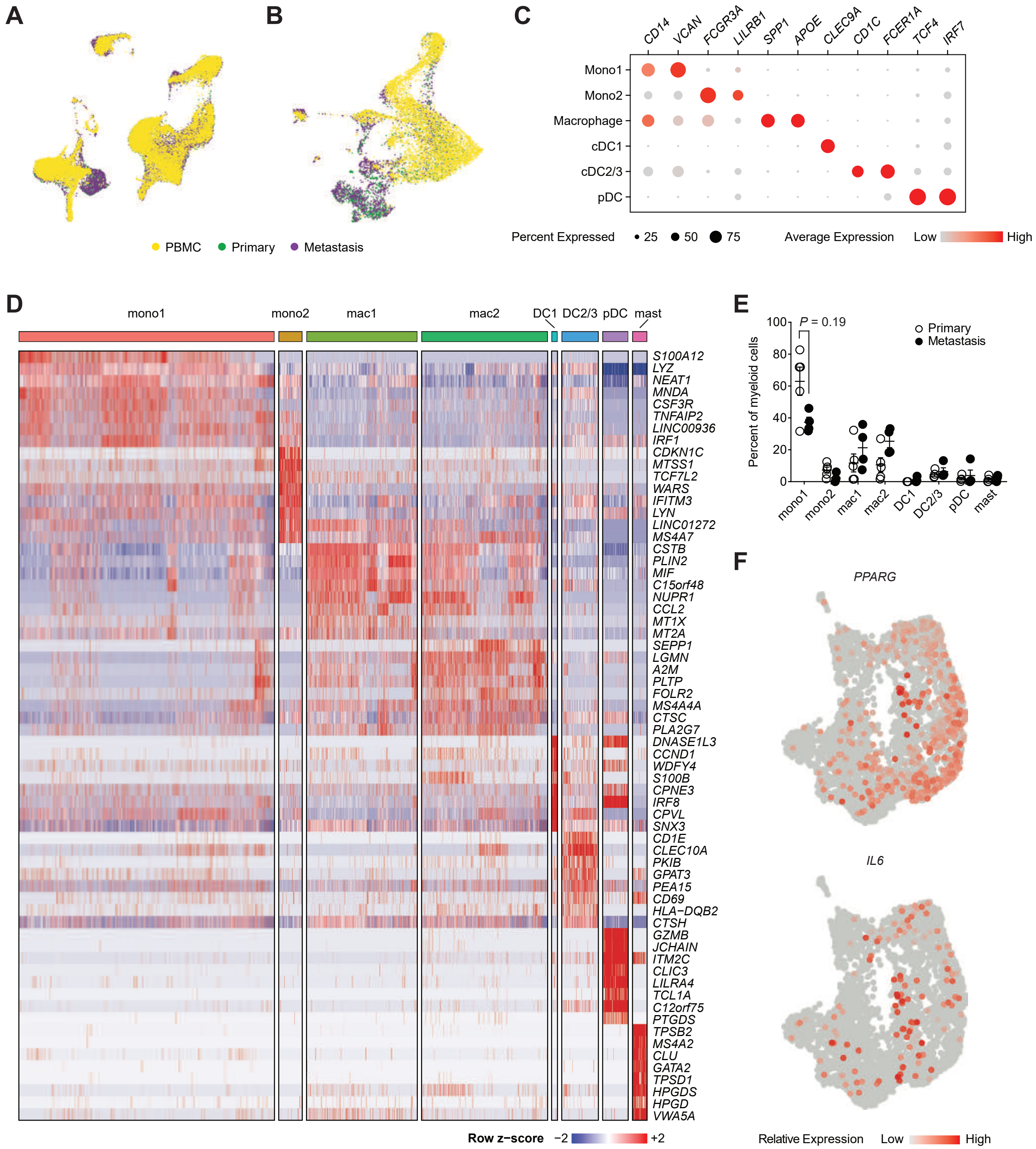

### Supplementary Figure S6

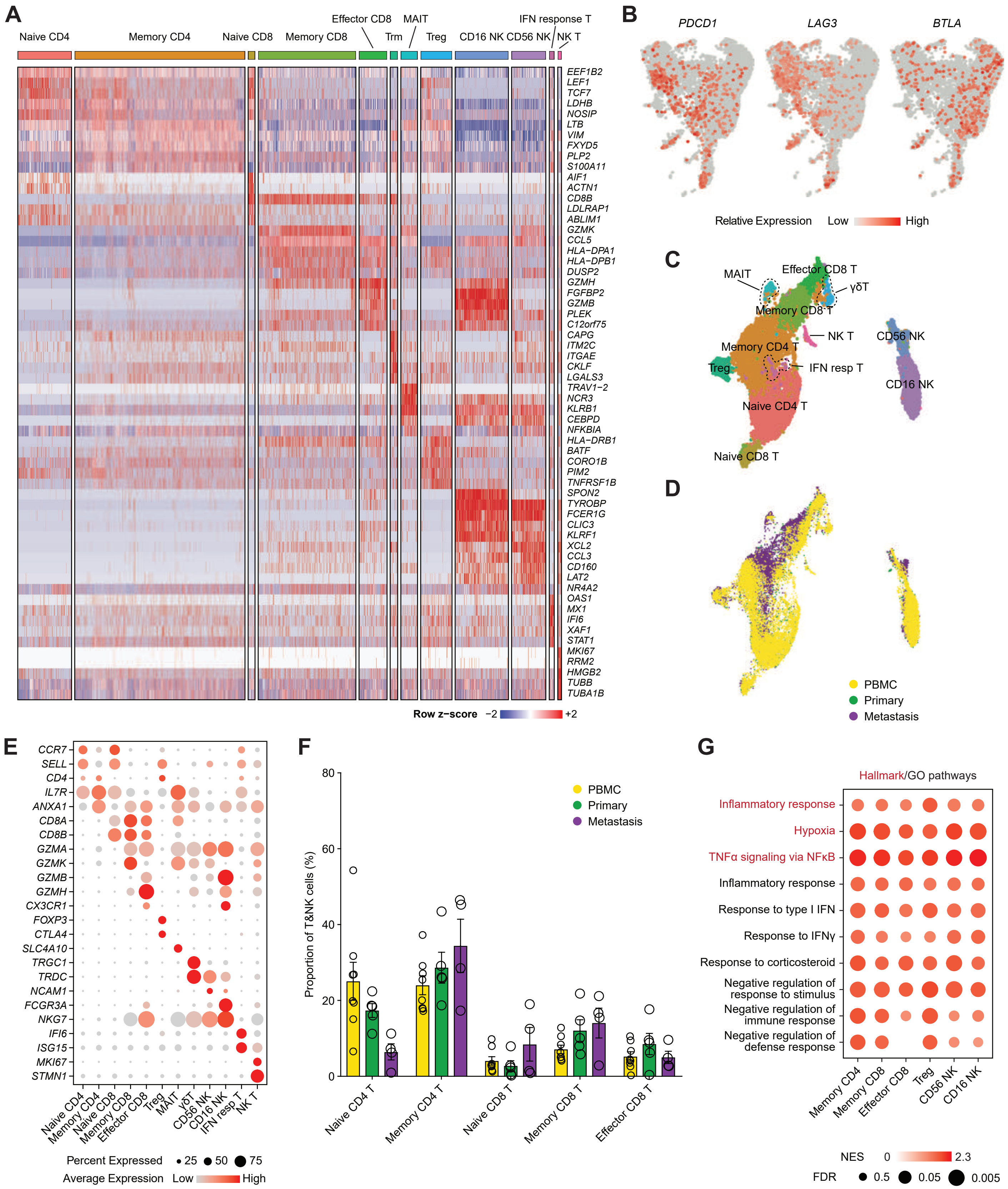

### Supplementary Figure S7

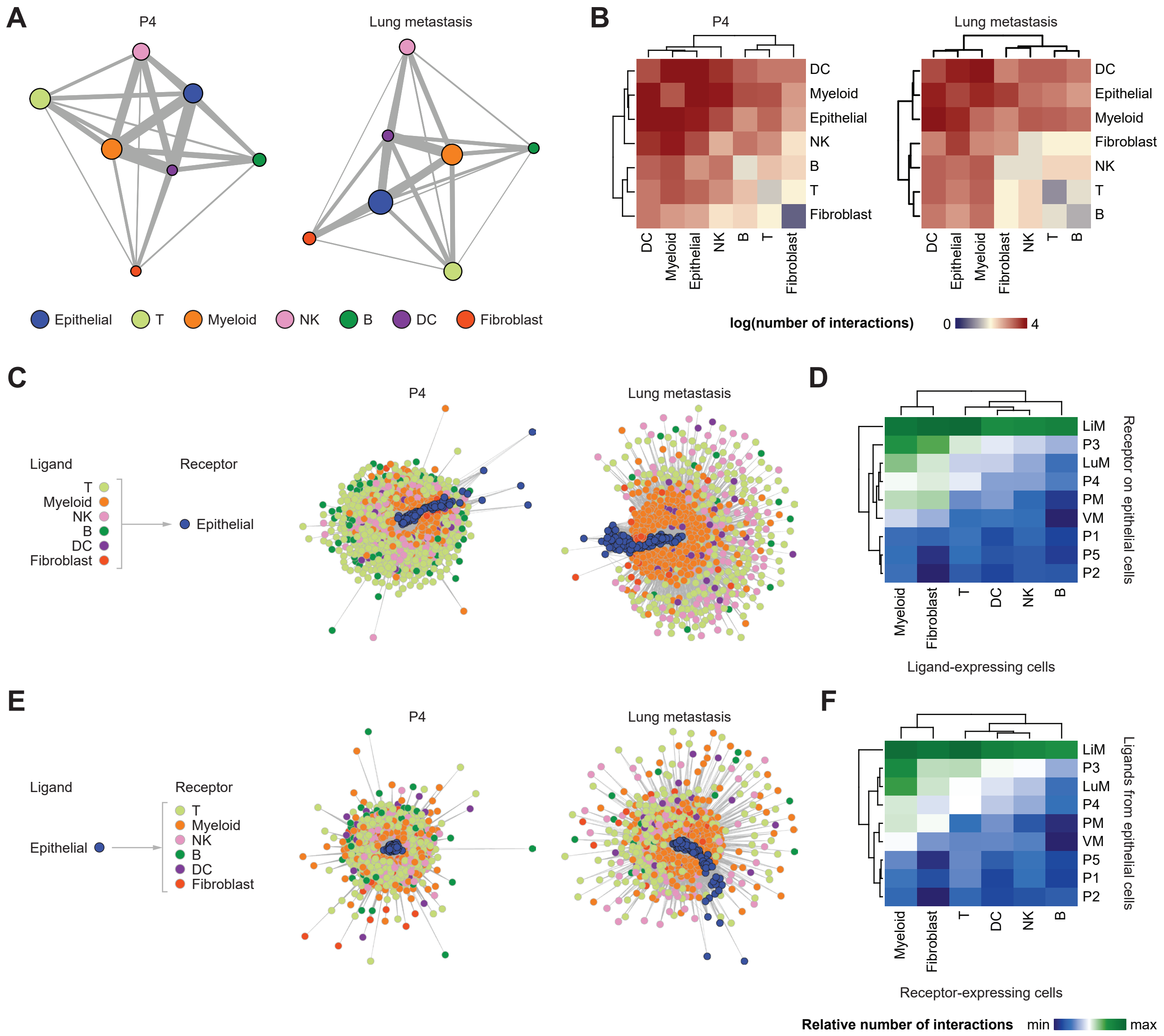

**Supplementary Table 1.** Total cell counts summarized by cell type and sample type (primary or metastasis).

| Cell Type | Primary | Metastasis | Total |
|---|---|---|---|
| **Epithelial** | 6,388 | 8,546 | 14,934 |
| **T** | 3,388 | 5,491 | 8,879 |
| **Myeloid** | 2,017 | 1,910 | 3,927 |
| **NK** | 788 | 868 | 1,656 |
| **B** | 297 | 1,036 | 1,333 |
| **DC** | 58 | 181 | 239 |
| **Endothelial** | 30 | 387 | 417 |
| **Fibroblast** | 41 | 294 | 335 |
| **Sum** | 13,007 | 18,713 | 31,720 |

**Supplementary Table 2.** Summary of epithelial sub-cluster cell counts by sample type.

| Cell Type | Primary | Metastatic | Total |
|---|---|---|---|
| **Epithelial 1** | 2,792 | 3,519 | 6,311 |
| **Epithelial 2** | 999 | 1,565 | 2,564 |
| **Epithelial 3** | 858 | 1,595 | 2,453 |
| **Epithelial 4** | 344 | 494 | 838 |
| **Lipid processing** | 399 | 129 | 528 |
| **Secretory** | 13 | 106 | 119 |
| **Sum** | 5,405 | 7,408 | 12,813 |

**Supplementary Table 3.** Results of molecular subtyping of all cells by Bailey, Collisson, and Moffitt classifiers. Numbers are organized by sample of origin and sample type.

| | Bailey | | | | | Collisson | | | | Moffitt | | |
|---|---|---|---|---|---|---|---|---|---|---|---|---|
| Sample | Pancreatic progenitor | Squamous | ADEX | Immunogenic | Sum | Classical | QM | Exocrine | Sum | Classical | Basal | Sum |
| P1 | 29 | 10 | 499 | 9 | 547 | 28 | 2 | 490 | 520 | 38 | 64 | 102 |
| P2 | 56 | 43 | 19 | 147 | 265 | 19 | 3 | 4 | 26 | 16 | 79 | 95 |
| P3 | 458 | 237 | 100 | 36 | 831 | 3,489 | 153 | 115 | 3,757 | 3,233 | 207 | 3,440 |
| P4 | 246 | 133 | 58 | 35 | 472 | 536 | 73 | 13 | 622 | 284 | 248 | 532 |
| P5 | 34 | 16 | 27 | 15 | 92 | 65 | 1 | 7 | 73 | 54 | 43 | 97 |
| VM | 698 | 55 | 7 | 2 | 762 | 1,432 | 44 | 9 | 1,485 | 1,283 | 28 | 1,311 |
| LiM | 356 | 1,582 | 216 | 567 | 2,721 | 247 | 1,013 | 49 | 1,309 | 1 | 4,325 | 4,326 |
| LuM | 64 | 1,056 | 35 | 12 | 1,167 | 826 | 409 | 8 | 1,243 | 104 | 995 | 1,099 |
| PM | 36 | 98 | 6 | 0 | 140 | 232 | 102 | 0 | 334 | 191 | 6 | 197 |
| Primary | 823 | 439 | 703 | 242 | 2,207 | 4,137 | 232 | 629 | 4,998 | 3,625 | 641 | 4,266 |
| Metastasis | 1,154 | 2,791 | 264 | 581 | 4,790 | 2,737 | 1,568 | 66 | 4,371 | 1,579 | 5,354 | 6,933 |
| Sum | 1,977 | 3,230 | 967 | 823 | 6,997 | 6,874 | 1,800 | 695 | 9,369 | 5,204 | 5,995 | 11,199 |

**Supplementary Table 4.** Results of molecular subtyping of epithelial cells by Bailey, Collisson, and Moffitt classifiers. Numbers are organized by sample of origin and sample type.

| Sample | Bailey | | | | | Collisson | | | | Moffitt | | |
|---|---|---|---|---|---|---|---|---|---|---|---|---|
| | Pancreatic progenitor | Squamous | ADEX | Immunogenic | Sum | Classical | QM | Exocrine | Sum | Classical | Basal | Sum |
| **P1** | 9 | 2 | 42 | 2 | 55 | 6 | 0 | 62 | 68 | 36 | 11 | 47 |
| **P2** | 3 | 0 | 7 | 0 | 10 | 8 | 0 | 2 | 10 | 15 | 2 | 17 |
| **P3** | 248 | 35 | 134 | 82 | 499 | 1,611 | 49 | 102 | 1,762 | 2,304 | 126 | 2,430 |
| **P4** | 142 | 27 | 11 | 4 | 184 | 324 | 24 | 6 | 354 | 274 | 95 | 369 |
| **P5** | 4 | 2 | 5 | 1 | 12 | 6 | 0 | 8 | 14 | 40 | 8 | 48 |
| **VM** | 624 | 0 | 25 | 29 | 678 | 487 | 2 | 6 | 495 | 1,092 | 23 | 1,115 |
| **LiM** | 126 | 1,232 | 45 | 4 | 1,407 | 10 | 516 | 6 | 532 | 2 | 2,199 | 2,201 |
| **LuM** | 22 | 1,033 | 58 | 28 | 1,141 | 114 | 352 | 10 | 476 | 93 | 1,520 | 1,613 |
| **PM** | 5 | 6 | 5 | 0 | 16 | 118 | 9 | 1 | 128 | 112 | 4 | 116 |
| **Primary** | 406 | 66 | 199 | 89 | 760 | 1,955 | 73 | 180 | 2,208 | 2,669 | 242 | 2,911 |
| **Metastasis** | 777 | 2,271 | 133 | 61 | 3,242 | 729 | 879 | 23 | 1,631 | 1,299 | 3,746 | 5,045 |
| **Sum** | 1,183 | 2,337 | 332 | 150 | 4,002 | 2,684 | 952 | 203 | 3,839 | 3,968 | 3,988 | 7,956 |

**Supplementary Table 5.** Summary of fibroblast subtypes by sample type.

| Cell Type | Primary | Metastatic | Total |
|---|---|---|---|
| apCAF | 29 | 72 | 101 |
| iCAF | 27 | 72 | 99 |
| myCAF | 14 | 182 | 196 |
| Sum | 70 | 326 | 396 |

**Supplementary Table 6.** Summary of myeloid cell subtypes by sample type.

| Cell Type | Primary | Metastatic | Total |
|---|---|---|---|
| **Monocyte 1** | 925 | 673 | 1,598 |
| **Monocyte 2** | 73 | 74 | 147 |
| **Macrophage 1** | 385 | 308 | 693 |
| **Macrophage 2** | 328 | 460 | 788 |
| **DC1** | 0 | 37 | 37 |
| **DC2/3** | 60 | 169 | 229 |
| **pDC** | 17 | 143 | 160 |
| **Mast** | 43 | 48 | 91 |
| **Sum** | 1,831 | 1,912 | 3,743 |

**Supplementary Table 7.** Summary of T and NK cell subtypes by sample type.

| Cell Type | Primary | Metastatic | Total |
|---|---|---|---|
| **Naïve CD4 T** | 611 | 373 | 984 |
| **Memory CD4 T** | 1,368 | 2,620 | 3,988 |
| **Naïve CD8 T** | 93 | 33 | 126 |
| **Memory CD8 T** | 381 | 1,406 | 1,787 |
| **Effector CD8 T** | 391 | 121 | 512 |
| **Trm** | 84 | 53 | 137 |
| **MAIT** | 53 | 252 | 305 |
| **Treg** | 196 | 377 | 573 |
| **CD16 NK** | 576 | 398 | 974 |
| **CD56 NK** | 169 | 456 | 625 |
| **IFN response T** | 22 | 61 | 83 |
| **NK T** | 14 | 49 | 63 |
| **Sum** | 3,958 | 6,199 | 10,157 |
